# Strong interactions between highly dynamic lamina-associated domains and the nuclear envelope stabilize the 3D architecture of Drosophila interphase chromatin

**DOI:** 10.1101/2022.01.28.478236

**Authors:** Igor S Tolokh, Nicholas Allen Kinney, Igor V Sharakhov, Alexey V Onufriev

## Abstract

**Background:** Interactions among topologically associating domains (TADs), and between the nuclear envelope (NE) and lamina-associated domains (LADs) are expected to shape various aspects of 3D chromatin structure and dynamics; however, relevant genome-wide experiments that may provide statistically significant conclusions remain difficult.

**Results:** We have developed a coarse-grained dynamical model of the *Drosophila melanogaster* nuclei at TAD resolution that explicitly accounts for four distinct epigenetic classes of TADs and LAD-NE interactions. The model is parameterized to reproduce the experimental Hi-C map of the wild type (WT) nuclei; it describes time evolution of the chromatin over the G1 phase of the interphase. Best agreement with the experiment is achieved when the simulations include an ensemble of nuclei, corresponding to the experimentally observed set of several possible mutual arrangements of chromosomal arms. The model is validated against multiple structural features of chromatin from several different experiments not used in model development, including those that describe changes in chromatin induced by lamin depletion. Predicted positioning of all LADs at the NE is highly dynamic – the same LAD can attach, detach and move far away from the NE multiple times during interphase. The probabilities of LADs to be in contact with the NE vary by an order of magnitude, despite all having the same affinity to the NE in the model. These probabilities are mostly determined by a highly variable local linear density of LADs along the genome which also has a strong effect on the predicted radial positioning of individual TADs. Higher probability of a TAD to be near NE is largely determined by a higher linear density of LADs surrounding this TAD. The distribution of LADs along the chromosome chains plays a notable role in maintaining a non-random average global structure of chromatin. Relatively high affinity of LADs to the NE in the WT nuclei substantially reduces sensitivity of the global radial chromatin distribution to variations in the strength of TAD-TAD interactions compared to the lamin depleted nuclei, where a 0.5 *kT* increase of cross-type TAD-TAD interactions doubles the chromatin density in the central nucleus region.

**Conclusions:** A dynamical model of the entire fruit fly genome makes multiple genome-wide predictions of biological interest. The distribution of LADs along the chromatin chains affects their probabilities to be in contact with the NE and radial positioning of highly mobile TADs, playing a notable role in creating a non-random average global structure of the chromatin. We conjecture that an important role of attractive LAD-NE interactions is to stabilize global chromatin structure against inevitable cell-to-cell variations in TAD-TAD interactions.

## Introduction

Interphase chromosomes are intricately folded and packaged inside the cell nuclei where they interact with nuclear bodies and remodeling factors [1, 2, 3, 4, 5]. This complex 3D organization of the genome plays a crucial role in regulation of gene expression, DNA replication, and DNA double-strand break repair [6, 7, 8, 9, 10, 11, 12, 13]. Chromosomes are spatially partitioned into sub-megabase topologically associating domains (TADs), which have emerged as the fundamental structural and functional units of chromatin organization in the eukaryotic cell nuclei, from fruit fly to human [1, 14, 15, 2]; [16, 17]. Interactions among TADs contribute to higher level, hierarchical genome organization [18]: multiple TADs within each chromatin compartment and multiple compartments within each chromosome territory. Boundaries of TADs are conserved among cell types and sometimes across species [14, 19, 20, 21, 22, 16, 23, 24, 25]; this feature makes TAD a natural structural unit of chromatin useful for modeling purposes and reasoning [26, 27], including in this work.

TADs interact among themselves forming dynamic, relatively short-living 3Dstructures, which segregate chromatin into mutually excluded active (type A) and inactive (type B) compartments [28, 18]. Averaged over time of the entire interphase, the effect of TAD-TAD interactions is seen in experimental Hi-C maps course-gained to TAD resolution [1, 14, 29, 26]. However, such Hi-C maps do not reveal how mobile are the 3D positions of each individual TAD. Live-cell imaging studies of interphase nuclei in different organisms showed that some chromatin loci are mobile [30, 31, 32, 33, 34, 35]. Also, studies suggested that constrains on the motion of chromatin loci can result from their interaction with structural elements such as nucleolus [36], or nuclear lamina (NL) which lines the inner surface of the nuclear envelope (NE) [37], or the nuclear pore complexes [31, 38, 39, 40, 41]. However, genome-wide analysis of nuclear dynamics of all “freely floating” and “tethered” chromatin loci has not been performed. DamID experiments have identified hundreds of genome regions that can “anchor” to the NL [42, 43, 37, 44, 45]. These genome regions that have specific affinity to the NL are called lamina-associated domains (LADs) [46, 47, 48, 49].

Positioning of LADs in mammalian cells [34, 50, 48, 47, 51] was shown to be stochastic: averaged across cells, a certain % of LADs is bound to the NE in a given cell, but the binding of specific LADs varies from cell to cell. Single-cell DamID studies demonstrate [34, 50] that LADs identified in a human cell population may in fact be located either at the nuclear periphery (30% of LADs) or in the nuclear interior in individual cells [34]. The LADs that contacted the NL were shown to move during interphase [34], but this movement is confined to a relatively narrow 1 *μ*m layer next to the NE. These observations raise several questions [49]. Is positioning of LADs only stochastic between cells (i.e., LADs have mostly stable positions within a cell, but highly variable positions between cells) or is it truly dynamic/mobile in each cell (i.e., individual LADs can change their positions relative to the NE significantly within a cell during interphase). If LADs are mobile, on what timescale? Also, do different LADs have different probabilities of being attached to the lamina?

States of transcription activation and repression are also linked to the positioning of chromatin with respect to the NE. LADs are typically repressive chromatin environment [42, 43, 45]. This observation supports the notion that the nuclear periphery is generally occupied by inactive chromatin [52]. TADs and LADs are similar in size: each is approximately 1 Mb in the human genome [14, 43] and approximately 90-100 kb in the *D. melanogaster* genome [46, 1, 15]. There are 1169 TADs and 412 LADs in the D. *melanogaster* genome [46, 1]. The role of LAD-NE interactions and their interplay with compartmentalization of TADs has been explored in *Drosophila* [35, 53] and mammals [54, 51, 55, 56, 57, 58, 59]. For example, depletion of lamins from *Drosophila* S2 cells leads to chromatin compaction and reduction in spatial segregation of the chromatin into active and inactive compartments [35]. Simulations of multiple copies of mouse chromosomes 1 and 2 [54] have disproved the role of lamina as the main driver of compartmentalization [47]. At the same time, attractions between heterochromatic TADs emerged as the main force of compartmentalization, while LAD-NE interactions are crucial for controlling the global spatial morphology of the nucleus [54, 56, 53]. However, a number of questions remains unanswered, related to sensitivity of the general organization principles with respect to global loss of chromatin-lamina interactions, which may occur in disease or senescence [51, 60, 61, 62, 63]. Also, the strength of TAD-TAD interactions can naturally vary during life of organisms. For example, the chromatin compartmentalization is weaker in embryonic cells and stronger in adult cells of *Anopheles* mosquitoes [64]. Given that attractive TAD-TAD interactions play a major role in compartmentalization, will the global chromatin architecture, such as its radial distribution, become more sensitive or less sensitive to the variations in these interactions upon lamin depletion? Will the cell-to-cell variability in the 3D chromatin organization increase or decrease upon disruption of LAD-NE interactions?

Understanding the principles and factors leading to the formation of non-random 3D genome organization and, ultimately, to the structure-function relationships in chromatin is prerequisite to understanding cell physiology [4, 65, 54]. However, relevant experiments that may provide statistically significant conclusions remain difficult. Computational models that faithfully reproduce available experimental data are indispensable for the 3D genome reconstruction problem (3D-GRP) based on DNA-proximity ligation data such as 5C, Hi-C and Pore-C, and they can generate valuable predictions and guide experiment [66, 67, 68, 69, 70, 71, 72, 73, 74, 75, 76, 77, 78, 79, 80, 26, 81, 82, 83, 84, 85, 86, 54, 51, 55, 87, 88, 89, 90, 91]. A particularly strong feature of computational models is that they can answer questions that may be very hard to address experimentally [92]. For example, tracing the movement of a given TAD or a few chromosomal loci upon detachment from the NE caused by a lamin depletion is possible experimentally [35, 93], but making statistically significant statements based on 1000s of TADs in the nucleus would be extremely laborious. Another example is obtaining time-resolved Hi-C maps for a synchronized cell population: possible, but difficult experimentally, with only a handful of studies so far [94, 95, 96, 97, 98] Models can generate testable hypotheses, the most promising of which can be checked experimentally.

Since simulating an entire eukaryotic genome on biologically meaningful timescales *at fully atomistic resolution* is still far out of reach [99], current practical models accept various levels of approximation determined by the balance of research goals and computational feasibility. A large class of recent computational models, which aim to understanding factors affecting 3D chromatin structure, employ the “beads-on-a-string” coarse-graining approach [67, 100, 101, 81, 76, 82, 51, 59, 89], borrowed from polymer physics models [102, 103]. For complex organisms, such as mammals, various further approximations are often made to reach the desired temporal or/and spatial resolution, such as considering only a small subset of the dozens of the original chromosomes [54] or even only one chromosome [51, 55, 58]. In that respect, modeling “simpler” nuclei [70, 89], in particular of higher eukaryotes, e.g., that of the well-studied fruit fly, with its only five major chromosome arms, offers a computational advantage that may translate into an ability to model the entire nucleus [76, 26, 82], which, in turn, may help to answer questions otherwise difficult to address. Consensus conclusions about chromatin organization that emerge from using models of different types applied to substantially different organisms are of value, as these conclusions hint at conservation of general principles across species.

Our work contributes to developing a qualitative and quantitative understanding of various aspects of 3D structure and dynamics of chromatin in *D. melanogaster* nuclei by constructing and employing a computational model to make various testable predictions. The model is trained to reproduce contact probabilities between TADs from the experimental Hi-C data [1, 26] and the average fraction of LADs near the NE [42], and is validated on multiple structural features of chromatin from several different experiments. It allows us to make multiple, biologically relevant predictions followed by a discussion about their possible biological significance. We have used the model to investigate sensitivity of the spatial organization of the interphase chromatin to the variation of the interactions among chromatin domains, clarifying the role of LAD-NE interactions in stabilization of the local and global chromatin structure. The model makes a genome-wide prediction of a highly dynamic nature of LAD positioning in fruit fly interphase nuclei – most of the LADs within a single nucleus attach to and detach from the NE multiple times, moving far away from the nuclear periphery on the time scale of the interphase. Previous experimental study [35] of LAD mobility in fruit fly was limited to only three LADs. Our model also predicts that the very different probabilities of individual LADs to be in contact with the NE are determined by the highly variable local linear density of LADs along the chromosomes which also affects the radial positioning of individual TAD, determining non-random average global structure of chromatin.

The dynamical model of the 3D chromatin organization in fruit fly proposed in this study offers a number of novel genome-wide insights, both biological and methodological, while also reinforcing robustness of some of the conclusions about chromatin organization previously made using models of mammalian nuclei. Combined with appropriate experimental data, our model can be used to make structurefunction predictions on genome-wide level [104].

## Methods

### Background and rationale for choosing model building blocks at TAD level

Our model is coarse-grained at the resolution of individual TADs. Numerous chromatin contacts within each TAD make the intra-TAD interactions effectively stronger than the inter-TAD ones. This difference allows one to consider the intra-TAD interactions as the major factors responsible for the shapes of TADs, while neglecting the effects of the inter-TAD and TAD-NE interaction on these shapes. We use these considerations and omit the details of the internal structure of TADs, considering them as the smallest building blocks of our model and using the shape-independent inter-TAD and TAD-NE interactions.

In *Drosophila melanogaster* nuclei each TAD compartmentalizes on average ~100 kb of chromatin [1]; there are four major epigenetic classes of TADs: Active, Null, PcG and HP1 [105, 1] (see Fig. 1 in the SI). Active TADs are defined by histone H3 modifications such as trimethylation of lysine 4 and 36 (H3K4me3 and H3K36me3). PcG TADs are enriched in Polycomb group proteins and histone mark H3K27me3. HP1 TADs are associated with classical heterochromatin marks such as H3K9me2 histone modification, heterochromatin proteins HP1 and Su(var)3-9. Null TADs lack known specific chromatin marks [1]. The differences in TAD-TAD interactions within each of these classes and between different classes are major factors responsible for the segregation of chromatin into active (type A) and inactive (type B) compartments [5, 54].

**Figure 1.**
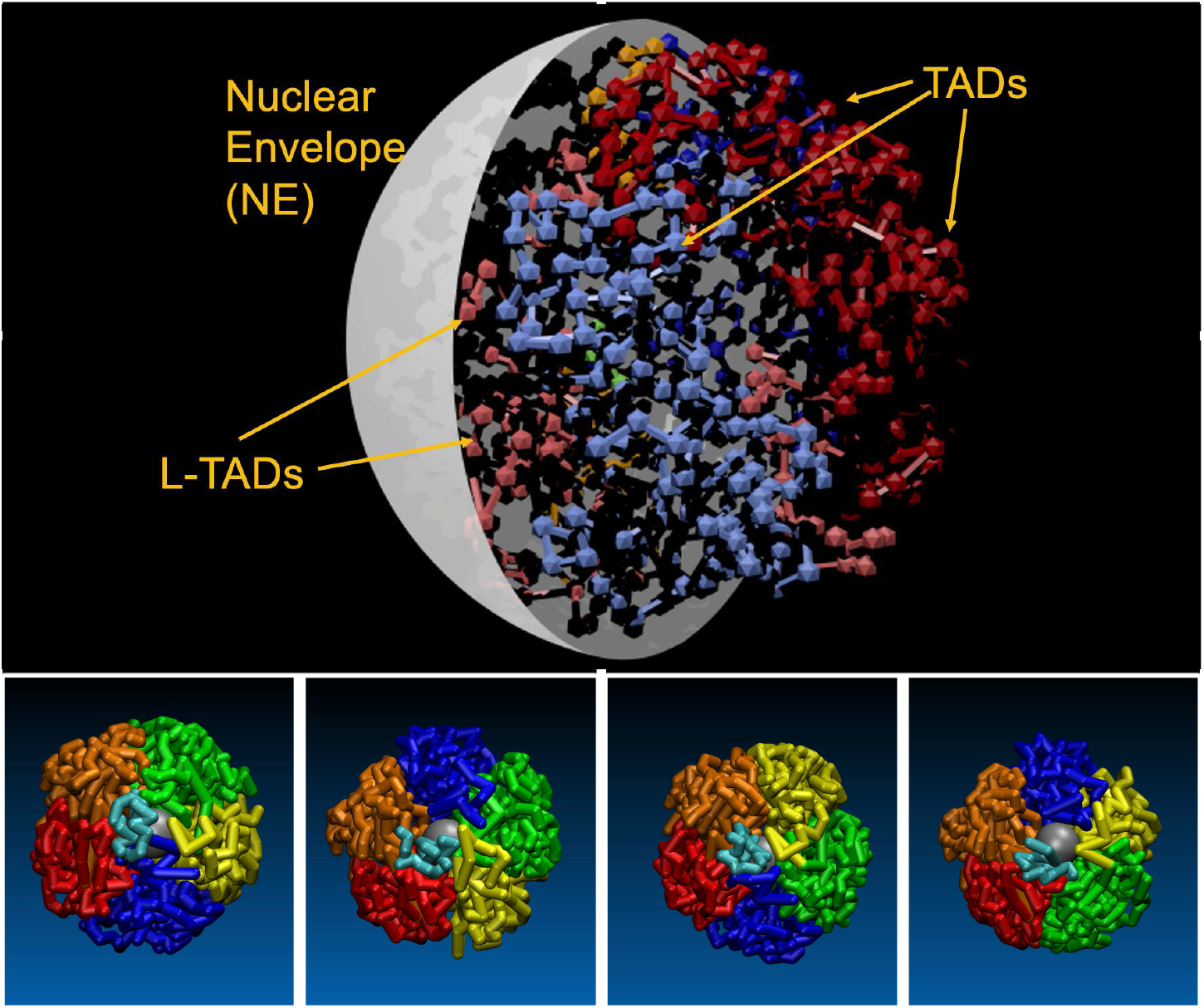
**Top:** A schematic showing the key elements of the model nucleus of *D. melanogaster.* **Bottom panels:** The four initial configurations (**bottom pannels**) of the *D. melanogaster* chromosome arms (2L (red), 2R (orange), 3L (yellow), 3R (green), X (blue) and 4 (cyan)) that serve as the starting points of the simulations described here. The nucleolus is shown as a grey sphere (on bottom panels). The arms are fully “territorial” at the beginning of each simulation. The initial configurations correspond to the different mutual arrangements of the chromosome arms (nucleus topologies) experimentally determined by Hochstrasser et al. [107]. **From left to right:** CIS-X6S, CIS-X7N, TRANS-X3S and TRANS-X4N nucleus topologies. Here, the “CIS” configuration refers to chromosome arrangement in which the two “R” arms (orange and green) or two “L” arms (red and yellow) of autosomes are next to each other in 3D space, while “TRANS” arrangement is complementary to it.

The four classes of TADs in fruit fly further refine the distinction between type A and type B compartments. Type A compartments are organized by Active class TADs and defined by early replication, proteins, and histone modifications involved in active transcription. Type B compartments are defined by late replication and modifications that silence genes. These compartments consist of transcriptionally silent TADs (PcG, HP1 and Null) and occupy a larger portion of the genome than type A compartments [1, 15, 106]

Even though our model is coarse-grained at TAD resolution and omits details of the internal structure of TADs, it provides a rather realistic description of the nuclear architecture by taking into account different mutual arrangements of chromosome arms (nucleus topologies) [107], different epigenetic classes of TADs [105, 1] and their interactions, and a proper distribution of LADs along chromosome chains [46].

### Model: chromosome representation

We consider *D. melanogaster* female interphase nuclei with a diploid set of four chromosomes (2, 3, 4 and X). Hi-C data suggest [1] that *D. melanogaster* genome (excluding pericentromeric constitutive heterochromatin (HET) and centromeric (CEN) regions) is organized into 1169 TADs. Since the same TADs in homologous chromosomes are almost always in close proximity to each other [108, 30, 26, 109, 110], we represent each pair of homologous TADs by a single spherical bead. In addition to these 1169 TAD-beads, we introduce 4 beads representing CEN regions of the four chromosomes and 6 beads representing HET domains.

Using the “beads-on-a-string” model [102, 111], four chains of the beads represent four paired homologous chromosomes (Chr). In Chr 2 and 3, the L- and R-arms are connected via three-bead HET-CEN-HET structure. Chr 4 begins from a CEN-HET bead pair, and Chr X ends with a HET-CEN two-bead structure. The Chr chains are placed inside of a spherical boundary (see Fig. 1, top panel) which represents the nuclear envelope (NE). We also consider a nucleolus which is modeled as a spherical bead of 0.333 *μ*m radius placed at half distance between the nucleus center and the NE [26].

### Bead size and mass

The mass *m_i_* of each bead, which represents a pair of homologous TADs, is related to the DNA sequence length *L_i_* (in bp) in the corresponding TAD and consists of the DNA mass (~660 Da/bp) and the mass of proteins associated with a nucleosome (132500 Da/nucleosome) [112, 113]. Using a 200 bp value for the nucleosome repeat length [114, 115, 116] one can get the following: *m_i_* = 2*L_i_*(132500/200 + 660).

We assume that the volume of each bead is proportional to its mass, and use the modified TAD radii determined in [26] (“hard radii”), scaled by a factor 1.254031 to reflect the double volume (two homologous TADs) of our beads.

See more details in the SI.

### Bead-bead interactions and bead types

See detailed description in the SI.

### Interactions in 268 specific pairs of remote loci

See detailed description in the SI.

### Arrangement of chromosome arms (nucleus topology) and preparation of their initial configurations

Several mutual arrangements of the chromosome chains that correspond to CIS and TRANS relative arrangements of the Chr 2 and 3 arms, and two most probable positions (“North”–“N” and “South”–“S”) of the Chr X have been observed experimentally [107] (see Fig. 1, bottom panels). The CIS arrangement refers to the one in which the R-arms (or the L-arms) of the autosomes are next to each other. The TRANS arrangement refers to the positions of the R- and L-arms of the autosomes next to each other. In the simulations, we use the following arrangements of the nucleus topology [107]: CIS-X6S, CIS-X7N, TRANS-X3S and TRANS-X4N. The CIS:TRANS arrangements ratio is assigned its experimental weight of 2, meaning that twice more replicas of CIS topologies are simulated. In addition, each of these 6 properly weighted arrangements has 3 replicas to account for 3 different nucleus sizes used. Thus, the entire simulated ensemble consists of 18 model nuclei.

To create initial configurations of the bead chains that represent chromosome (Chr) X, L- and R-arms of chromosomes (Chrs) 2 and 3, we generated them as separate linear strings of beads parallel to z-axis (the axis of centromere-to-telomere chromosome polarization). Each chain is initially separately restrained by two planes intersecting along the z-axis with the dihedral angle 72°. To collapse initially linear chain of beads into the nucleus sphere, a harmonic interaction between the nucleus center and each bead is applied using (see Eq. 3 in the SI) with the effective equilibrium length 0.99 *μ*m and harmonic spring parameter *κ* = 0.5 *kT*. A subsequent Langevin dynamics of the restrained chains over 10^5^ 1.36 ns time steps produced collapsed fractal-like Chr-arm configurations [111]. This harmonic interaction is removed from all subsequent simulation stages. The L- and R-arms of Chrs 2 and 3 are then connected through the CEN beads, Chr 4 is added at the “North” pole, and large restraining sphere is placed around the chromosomes. A set of short Langevin dynamics simulations with a decreasing radius of the restraining sphere brings the size of the model nuclei to their specified final values. The resulting configurations of all four chromosomes are presented in Fig. 1 and are used as the initial configurations for the simulations of the interphase chromatin dynamics.

### Nuclear envelope and its interactions with chromatin chains

We model the NE as a spherical boundary that restrains the motion of the chromosomes and can attract LADs [37, 46, 47, 48]. Following [82], we map positions of 412 LADs (a median size of LADs is about 90 kb [46]) onto the chains of 1169 TADs. If TAD contains LAD, then the corresponding bead can attractively interact with the NE. After mapping, we have determined 350 TADs that contain LADs (L-TADs). We describe all L-TAD–NE (LAD-NE) attractive interactions by the LJ-cos potential (see Eq. 4 in the SI) with a single well depth parameter *\*_L_*. This is a simplification of the L-TAD-NE interactions (eliminating of the dependence of these interactions on the LAD length in L-TAD) which allows us to investigate what other factors can affect L-TAD-NE contact probabilities and radial distributions.

Fraction of LADs at the NE is calculated as the average (over the described above ensemble of 18 nuclei) fraction of L-TAD beads within (position of bead centers) 0.09 *μ*m layer (average bead radius) at the NE.

To analyze the mobility of LADs we compute the probabilities of L-TADs (the centers of L-TAD bead) to be within a very thin, 0.2 *μ*m layer immediately adjacent to the NE. The thickness of this layer roughly corresponds to the average diameter of beads in our model. When L-TAD is in this layer we define it as being in contact with the NE.

Interactions of the NE with TADs not containing LADs are described by the purely repulsive potential (see Eq. 3 in the SI).

### Lamin Mutant model

We refer to the lamin depleted or lamin knock-down nuclei as “Lamin mutant” nuclei. Our Lamin mutant nucleus model has the L-TAD–NE affinity parameter *\*_L_* reduced to essentially zero (0.1 *kT*, see a more detailed description in the SI). All other interaction parameters are unchanged from the wild-type (WT) model. The Hi-C map for Lamin mutant model is calculated using long “11 hours (hrs)” trajectories (see “Matching simulation time to biological time” subsection in “Methods”.)

### Experimental chromatin density profiles

The results of the simulations are compared with the experimental chromatin density data [117], briefly described in the SI.

### Simulation of chromatin dynamics

Langevin dynamics simulations have been performed using ESPResSo 3.3.1 package [118] by solving the Langevin equation of motion:

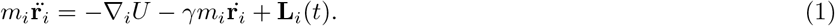

Here, **r**_*i*_ is the position of bead *i* with mass *m_i_*, *U* is the potential energy of the system. The last two terms describe the interaction with the solvent: a velocity dependent friction force, characterised by the parameter *γ*, and a random Gaussian white noise force **L**_*i*_. The integration time step is *t_step_* = 0.01*τ*, where 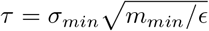 is the LJ time scale [103, 119]. Using *∊* = 3 *kT* we obtain *τ* = 136 ns and, accordingly, *t_step_* = 1.36 ns, which was used for all our simulations. For the simulations used to narrow down the range of the interactions parameters, we generated 40 × 10^6^ time step trajectories with *γ* = 1/*τ*. For the production simulations with the selected WT and Lamin mutant interaction parameter sets, we generated 400 × 10^6^ time steps trajectories using *γ* = 0.01/*τ*. This much smaller *γ* allows one to effectively speed up the simulations (~ 20 times), which is one of the key benefits [120] of the implicit solvent approach used here. Also, in this regime of small friction the bead inertia and, accordingly, the difference in bead masses may become important.

### Contact probability (Hi-C) map

The TAD-TAD contact probability map (Hi-C map) of the model is defined as the average of Hi-C maps (1169×1169) calculated over 18 trajectories of the ensemble of 18 model nuclei of 3 different sizes and 4 mutual arrangements of the chromosomes. (See a more detailed description in the SI).

The final Hi-C maps for the developed WT and Lamin mutant models are calculate using long “11 hrs” trajectories (see the following subsection).

### Matching simulation time to biological time

To relate the simulation timescale with the experiment [121], that is to map the simulation time onto real biological time, we compared [82] a diffusive motion of model beads with the experimental interphase chromatin diffusion, and used the match to estimate the scaling factor λ that converts the simulation time to real biological time. We calculated time dependencies of distance *R_i_*(*t*) between bead *i* and the nucleus center for 9 randomly selected beads, which do not contain LADs. The dependence of the mean squared displacement (MSD) on the time interval Δ*t*, 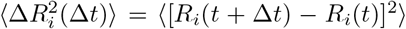, was calculated for 3 different values of the friction parameter *γ* (0.01/*τ*, 0.1/*τ* and 1/*τ*). The averaging was performed over 9 selected beads along the 18 trajectories. We fit the first 300 ·10^3^ time steps of each of the three curves 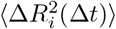 (see Fig. 3 in the SI) with the following equation for sub-diffusive motion of chromosomal loci [121]:

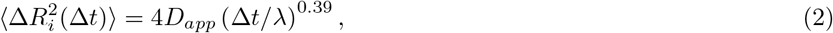

where *D_app_* is the apparent diffusion coefficient. This equation (with λ = 1 and Δ*t* in seconds) describes the experimentally observed diffusion of chromosomal loci over the time periods ranging from 1 to 10^3^ s [121]. We obtain reasonable fits of the simulation derived MSDs with 4*D_app_* = 0.061 *μ*m^2^ and λ = 10^4^ for *γ* = 0.01/*τ*, λ = 2.8 · 10^4^ for *γ* = 0.1/*τ*, and λ = 20 · 10^4^ for *γ* = 1/*τ* (for Δ*t* in the time steps). These values of λ give us the number of simulation time steps that corresponds to 1 sec of a real biological time. The scaling translates the 40 × 10^6^ time step simulations with *γ* = 1/*τ* into 3 minutes (min) of real nucleus time, and long production 400 × 10^6^ time step simulations with *γ* = 0.01/τ as corresponding to 667 min (11 hrs) of real time.

The Hi-C map calculated using long “11 hrs” trajectories for the WT parameter set has Pearson’s correlation coefficient with the experimental Hi-C map 0.954, close to 0.956 value obtained using “3 min” trajectories. These values suggest that reducing friction coefficient *γ* to 0.01 of its original 1/*τ* value used at stage of model parameters development, does not affect much of the local chromatin structure. A small deterioration of the original structure is expected due to time evolution and decay of the initial configurations.

Within the model, *t* = 0 corresponds to a point in the very beginning of G1 phase of the cell cycle, when all of the chromosomes are fully de-condensed.

### Model development

Our model consists of 1179 “soft” beads that represent homologous pairs of TADs (1169 beads), pericentromeric constitutive heterochromatin domains (6 HET beads) and centromeric chromatin domains (4 CEN beads) (see “Methods”). The beads representing TADs are split into four types which correspond to four major epigenetic TAD classes: Active, Null, PcG and HP1 [1]. The beads are combined into four chains of homologous chromosomes (Chrs 2, 3, 4 and X) using “beads-on-a-string” model [100, 81, 67, 101]. Nucleolus is modeled as an additional constrained bead [26].

The nuclear envelope (NE), a spherical boundary surrounding the chromosome chains (see Fig. 1), constrains the motion of the beads within the nucleus by repulsive interactions. At the same time, the NE can attract L-TADs (TADs that contain LADs) [42, 46]) representing LAD-NE interactions.

Compared to the previous fruit fly interphase chromatin model [82], and models developed for mammalian nuclei Refs. [122, 54, 51, 55], our model utilizes the attractive interactions between non-bonded beads. We assume that this presumably protein mediated TAD-TAD attraction [123, 124] is a TAD-class dependent [122], since different combinations of proteins are bound to different epigenetic classes of TADs [105] with their characteristic histone modifications.

Experimentally observed compartmentalization of chromatin [28] into active euchromatin (type A compartments) and more densely packed inactive heterochromatin (type B compartments) allows us to assume [54, 122] at least three types of TAD-TAD interactions: A-A, B-B and A-B. To allow the compartmentalization, some general relations (the so-called Flory-Huggins rule [125]) between these three types of TAD-TAD interactions have to be considered.

We also introduce a set of “specific” non-bonded TAD-TAD interactions for 268 TAD (bead) pairs which form “long-range” contacts with the increased probabilities [1]. These attractive interactions are described by a set of effective potentials with the well depths determined from the experimental contact probabilities for the corresponding TAD pairs (see “Methods” in the SI).

### Interaction parameters of the models

The attractive protein mediated interactions between TADs can spread over a several *kT* range of energies [126, 54].

To determine “optimum” bead-bead and L-TAD–NE interaction parameters of the model interphase nuclei we use simultaneously the following three major criteria (selection rules):

1. Maximum possible Pearson’s correlation coefficient between model derived TAD-TAD contact probability map (model Hi-C map) and the experimental WT Hi-C map [1], reduced to the TAD-TAD contact probabilities [26]);
2. The fraction of LADs (LAD containing beads, L-TADs) which are in contact with the NE matches the experiment for the WT nuclei: 25% [42]);
3. The commonly used condition for the chromatin compartmentalization is satisfied. This is the so-called Flory-Huggins rule [125]) – the strength of the interactions between different (A and B) unit types should satisfy the phase separation criterion: interaction A-B < (A-A + B-B)/2.

Three stages of model development are used (see the description in the SI, “Model development”).

The final optimum WT set of the interaction parameters is shown in the second column of the Table 1 below. We use this set for 10x longer simulations reported in the next section. The resulting chromatin density profiles will be compared with the available experimental data presented in [117]. We will also investigate how sensitive is the chromatin structure to the deviations in the parameters.

**Table 1.**
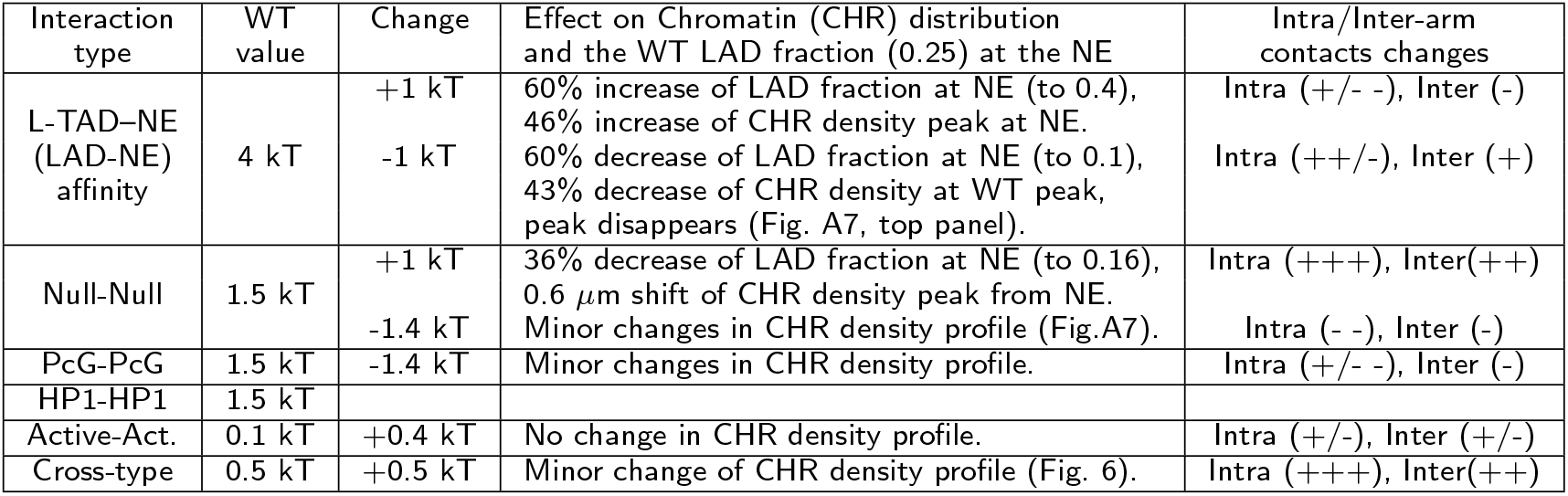
Wild-type (WT) model TAD interaction parameters and the effects of their variations on Chromatin (CHR) distribution and Chromosome arms contacts. Single “+” (“-”) denotes a small increase (decrease) in the corresponding contact probability. “++” (“- -”) denotes a moderate increase (decrease), while “+++” denotes a strong increase. ‘+/-” denotes increase and decrease.

## Results

### Dynamic model of chromosomes in fruit-fly at TAD resolution

We have developed a coarse-grained “beads-on-a-string” model of *D. melanogaster* female interphase nuclei at TAD resolution (the average TAD size is ~100 kb), where a bead represents a pair of homologous TADs in paired homologous chromosomes. Four main types of beads corresponding to four major epigenetic classes of TADs [1] – Active, Null, PcG and HP1 – are used. The arrangements of the chromosome chains (nucleus topologies) in the model nuclei correspond to the experimentally observed CIS and TRANS mutual arrangements of the L- and R-chromosome arms and the position of X chromosome [107], see Fig. 1 in “Methods”. Unless otherwise specified, the results are averages over the ensemble of all different chromosome topologies and nucleus sizes (18 systems in total, see “Methods”).

The main capabilities of the model are illustrated in Fig. 2, where we present the snapshots of the temporal evolution of the nucleus along the production trajectory, corresponding to about 11 hrs of biological time. The initial (*t* = 0) and the following (at *t* = 30 min, 3 hrs and 11 hrs) full ensemble averaged Hi-C maps are shown as well. The snapshots and the maps show the expected small decay [82] of the perfect original chromosome territories and local structures: the probabilities of TAD-TAD contacts within each chromosome arm slightly decrease with time (by about 5– 10%), although there are regions of contacts where these probabilities increase (see the difference maps in Fig. 5 in the SI). The presented configurations and the maps demonstrate that reasonably distinct territories still exist 11 hrs into the interphase, providing the first “sanity check” of our model. The over-all conclusion from the bottom panel of Fig. 2 (see also the difference maps in Fig. 5 in the SI) is that the Hi-C map of a fruit fly nucleus at TAD resolution experiences only relatively small changes at the time-scale of the interphase. We refrain from a more detailed analysis of these changes here, as our focus is the role of the NE.

**Figure 2.**
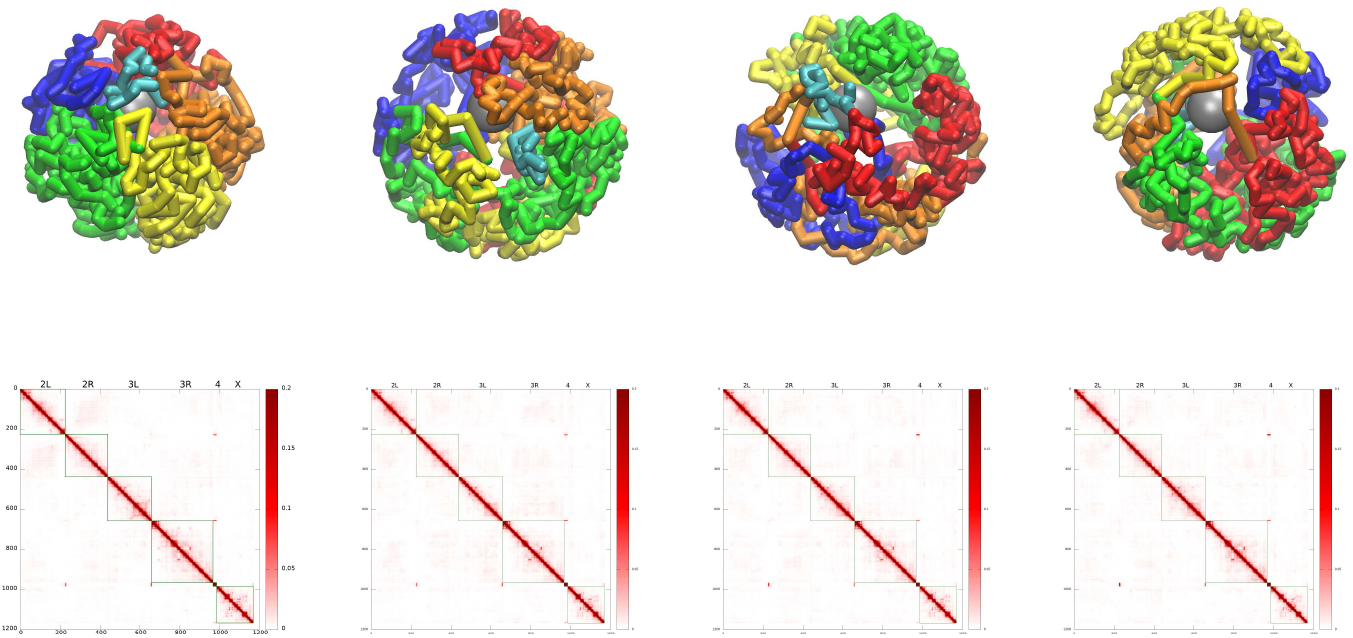
The temporal evolution of the WT chromatin configurations (**top row**), and the model derived TAD-TAD contact probability (Hi-C) maps (**bottom row**). **From left to right:** The model chromatin configurations at t=0 min (starting configuration), and at time points corresponding to 30 min, 3 hrs and 11 hrs. The configurations shown correspond to TRANS-X3S nucleus topology (see Fig. 1), used as an example. The Hi-C maps (bottom row) are each averaged over 5 min (3 · 10^6^ time steps) time intervals, and over the ensemble of 18 independent trajectories of the model nucleus, generated with the WT parameter set. The averaging includes all of the starting topologies, see “Methods”.

**Figure 3.**
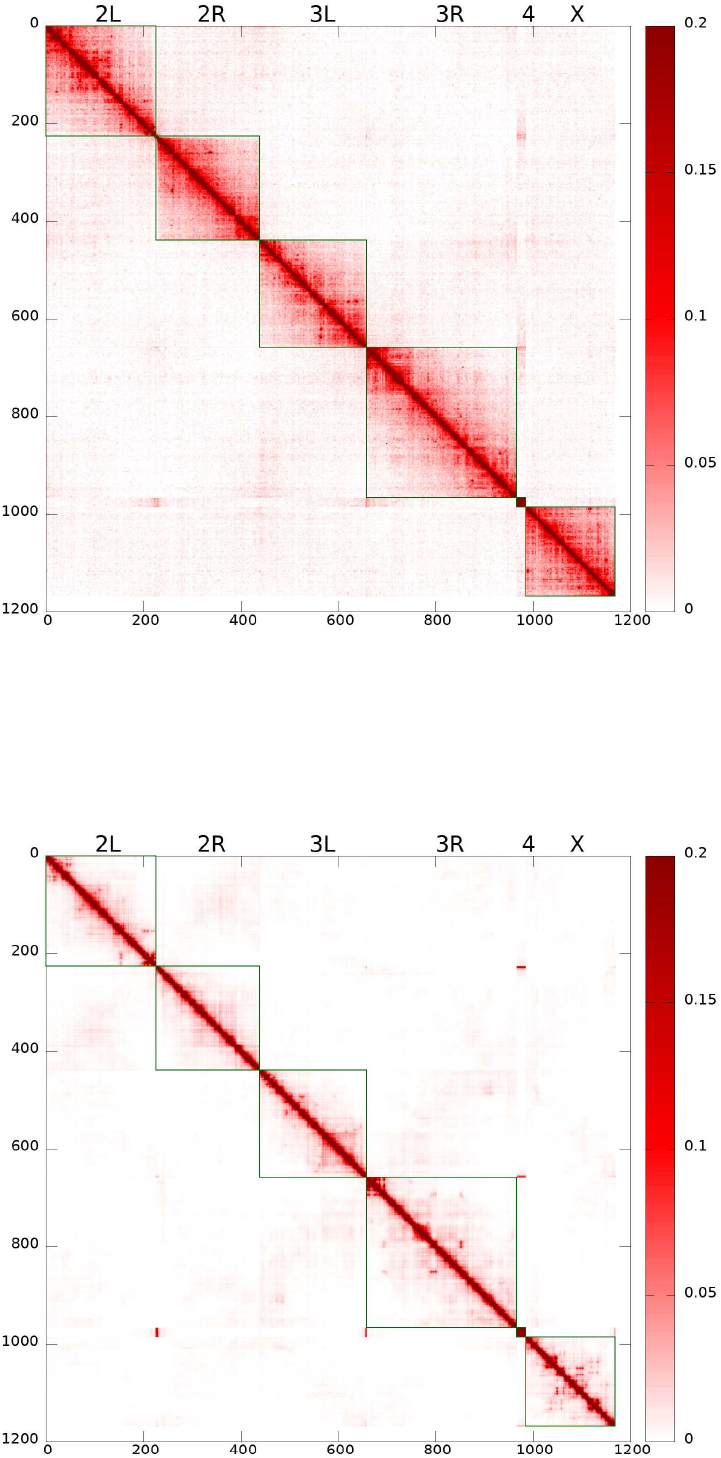
**Top panel:** Experimental Hi-C map for TAD-TAD contact probabilities in the WT *D. melanogaster* embryonic nuclei [26] (original data for embryonic nuclei from Ref. [1]). **Bottom panel:** Model derived TAD-TAD contact probability (Hi-C) map corresponding to young (0 – 3 min) nuclei. This Hi-C map reproduces many features expected from experiment: increased interactions within chromosomal arms, long-range chromatin contacts visible as bright spots, genome compartmentalization manifested as plaid-patterns of TAD-TAD contacts, and the Rabl-like configuration represented by interactions between chromosomal arms (e.g., 2L and 2R) as “wings” stretched perpendicular to the main diagonal. Pearson’s correlation coefficient with the experimental Hi-C map is 0.956.

**Figure 4.**
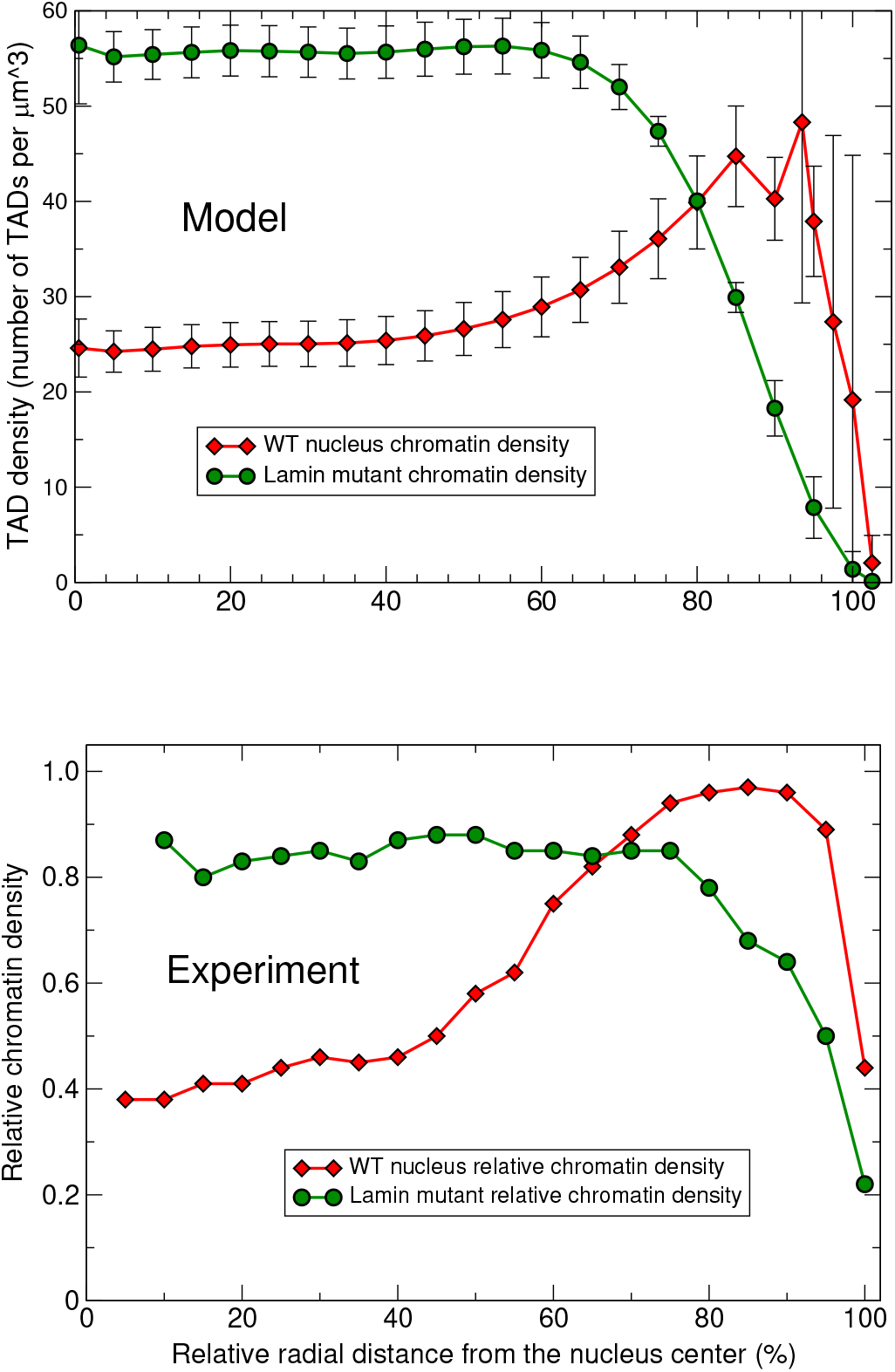
Radial chromatin density distributions in fruit fly nucelus. **Top panel:** Model WT nuclei (red curve, diamonds) and the Lamin mutant model nuclei (green curve, circles). The error bars are the standard deviations of the mean values for 18 nuclei. **Bottom panel:** Experimental WT nuclei (red curve, diamonds) and the Lamin mutant nuclei (green curve, circles) [Adapted from Supplementary Data, Figure S3 (Group 1) of Ref. [117]].

**Figure 5.**
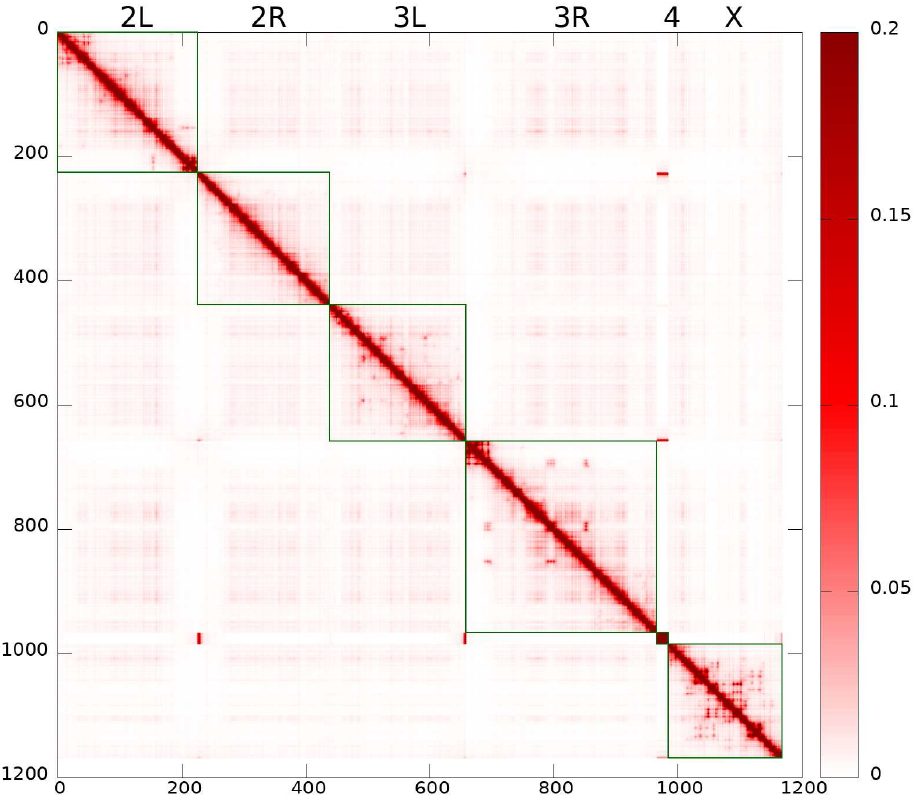
Model derived Lamin mutant TAD-TAD contact probability (Hi-C) map. Pearson’s correlation coefficient relative to the corresponding WT model Hi-C map is 0.9989, suggesting a high over-all similarity between the WT and Lamin mutant contact maps.

The model is trained to agree with the experimental Hi-C map, Fig. 3 (top panel) [1, 26], and lamin-DamID data [42] for the WT nuclei, and to satisfy the general polymer physics restrictions imposed on the strengths of the attractive interactions between beads of different types. The model derived TAD-TAD contact probability map, i.e., the model Hi-C map, Fig. 3 (bottom panel), has been calculated for the ensemble of “young” nuclei, corresponding to about 3 min of real biological time.

The selected WT parameter set provides a reasonably accurate reproduction of the experimental Hi-C map (Pearson’s correlation coefficient is 0.956). The apparent difference in the intensity of inter-arm and intra-arm TAD-TAD contacts between the experimental and model Hi-C maps can be explained by a more noisy experimental data and a rather short (~3 min) effective biological time for the model nuclei used to collect the data.

The model is validated against multiple (ten) structural features of chromatin from several different experiments, which were not used for its training, see below. The model is capable of predicting the evolution of the 3D chromatin architecture (see Fig. 2) on time scales up to 11 hrs, which is enough to approximate the duration of the G1 phase of the fruit fly nuclei interphase [127, 128, 129, 130, 30]. The Hi-C map calculated using “11 hrs” trajectories for the WT parameter set has Pearson’s correlation coefficient with the experimental [1] Hi-C map 0.954.‘

### WT nucleus vs. Lamin mutant nucleus

To assess the role of the NE in organization of the chromatin architecture, a Lamin depleted nucleus model (simply referred to as “Lamin mutant”) has been created by reducing the LAD-NE affinity from relatively strong 4 *kT* WT value to essentially zero, see“Methods”. The rest of the model parameters were kept the same.

Two model derived WT and Lamin mutant radial TAD density distributions are shown in Fig. 4 (top panel). These distributions are in reasonable agreement with recently reported experimental chromatin density distributions [117] shown in Fig. 4 (bottom panel). In particular, the model faithfully reproduces non-trivial nuances of all of the experimental distributions reported in Ref. [117], particularly the “flatness” of the of density profile away from the NE for Lamin mutant (in experimentally seen distributions), as opposed to, *e.g.,* continued growth of the density toward the center of the nucleus that can occur if the model parameters deviate from their optimal values, see below. This agreement provides a strong validation to the model, independent of the experimental data (WT Hi-C map) used for its training.

Comparing the WT and Lamin mutant chromatin distributions, Fig. 4, one can see a substantial shift of the chromatin density toward the NE in the WT nuclei. LAD-NE attractive interactions in the WT nuclei transform a more compact globulelike distribution in the Lamin mutants, with a very low chromatin density near the NE, into a more extended distribution, with a chromatin density peak near the NE and substantially reduced chromatin density in the central nucleus region. Similar effects of increased chromatin compaction and its movement away from the NE upon reduction of the LAD-NE attraction strength were previously observed in the experiments with *Drosophila* S2 cells [35], in the models of single human chromosomes [51, 55, 58] and in the model of several mouse chromosomes [54].

Comparing the model derived Hi-C maps for the Lamin mutant (Fig. 5) with that of the WT nuclei, our first conclusion is that the two are quite similar: the Pearson’s correlation coefficient between the two is 0.9989. Which means that, by and large, the significant global re-arrangements of the chromatin upon abrogation of the LAD-NE attraction, Fig. 4, have relatively little effect on the over-all structure TAD-TAD contacts, as revealed by the Hi-C map. The relatively subtle differences between the two Hi-C maps are best revealed in the difference map, see SI, Fig. 8. One can see that the small (within 10%) changes in the contact probabilities are in agreement with the changes in the chromatin density distribution upon transition to the Lamin mutant seen in Fig. 4. A more compact and dense chromatin in the Lamin mutant has slightly higher TAD-TAD contact probabilities, both within the chromosome arms and between the arms.

Also, the specific TAD-TAD contacts produce more intensive spots on the Lamin mutant map (Fig. 5) compared to the WT model Hi-C map (see the difference map in the SI, Fig. 8).

The model also reproduces key qualitative result from Ref. [35] that lamin depletion enhances interactions between active and inactive chromatin, impairing spatial segregation of active and inactive compartments. Our models provide a quantitative estimate for the increase of contacts between Null and Active TADs in the Lamin mutant relative to the WT nuclei (see the sums of the selected contact probabilities for each Null TAD with Active TADs in Fig. 11 in the SI). The averaged relative increase is 22%. This number is a testable prediction; the predicted increase is yet another validation of the model, independent of the data sets used to build the model.

### NE has a dual role, acting as an “attractive enclosure”

Comparing the model derived Lamin mutant and WT chromatin distributions (Fig. 4, top panel) one can see that the main consequence of the almost complete elimination of the LAD-NE affinity is a significant “global” change in the chromatin density profile. The peak of the WT chromatin density at about 1.9 *μ*m from the nucleus center (at about 94% of the relative distance from the center) disappears, and the chromatin is shifted away from the NE. The density in the nearest to the NE layer of TADs drops 6 times compared to the WT value, while the density in the central region of the nucleus doubles, making the resulting chromatin distribution more compact. A similar general conclusion about the ability of LAD-NE interactions to re-distribute large blocks of chromatin was reached in the context of models of single human chromosomes interacting with the NE [51, 58]. We note that the respective parameter regimes that lead to this common conclusion, *e.g.*, the ratios of TAD-TAD to LAD-NE interaction strengths, may be quite different from ours. On the other hand, it is reassuring that certain general principles of chromatin organization appear to be robust to model details.

What we have found unexpected is how quickly – on the minute time-scale – the more compact chromatin structure of the Lamin mutants deteriorates once the entire confining NE (not just Lamins) is removed completely, see Fig. 6 in the SI. These combined results suggests a dual role for the NE. It is not about mere confinement of chromatin; but also it is not simply “keeping interphase chromosomes slightly stretched” [35]: rather, the NE has a dual role, acting as an “attractive enclosure”, which can both redistribute the chromatin, shifting most of it from the interior to the nuclear periphery due to the LAD-NE attraction in the WT nuclei, and confine it, preventing from decondensation in Lamin mutants, in the absence of the LAD-NE attraction. Without the NE, the chromatin will not shrink as it does in Lamin mutants. Possible biological implications of these findings are touched upon in “Discussion”.

**Figure 6.**
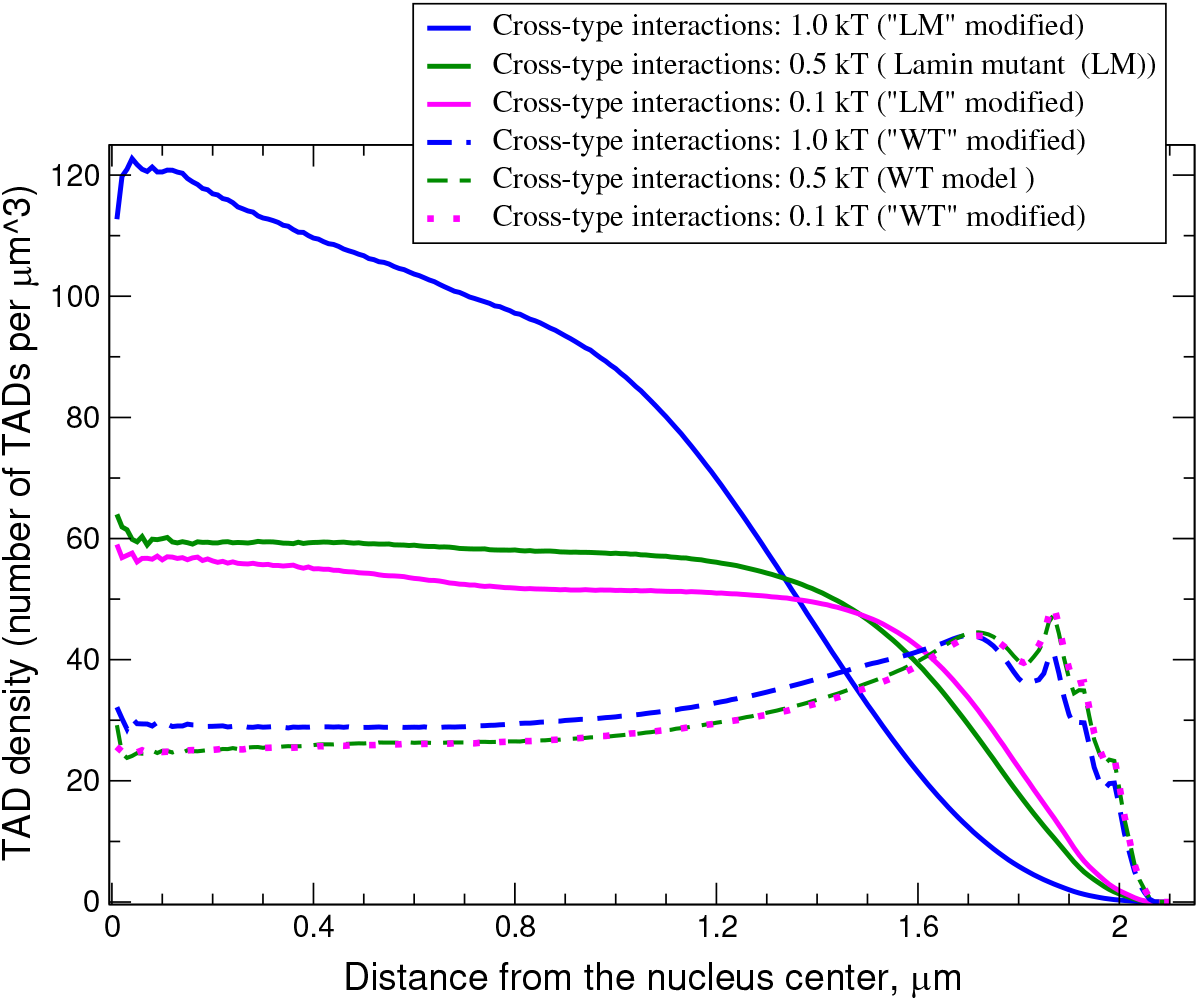
Chromatin density distributions in the WT and Lamin mutant nuclei models with modified levels of cross-type TAD-TAD attractive interactions (parameter *∊*_*A,B*_), both between A and B types of beads and between different types of B beads (e.g., Null-PcG interaction).

### 3D chromatin architecture is sensitive to variations in the interactions of its key elements

To investigate how different interaction types affect the chromatin properties we have simulated the nuclei with the interaction parameters deviating from the WT and the corresponding Lamin mutant set values.

#### Effect of LAD-NE affinity variation

One of the question which clarifies the role of the NE is how the deviation of the LAD-NE affinity from its WT value (4 *kT*) affects the chromatin distribution? In Fig. 7 in the SI (top panel), the radial distributions of TADs in the model at different levels of LAD-NE affinity (from 0.1 *kT* in Lamin mutant to 5 *kT*) are presented. One can see that the increase of LAD-NE affinity by 0.5 *kT* from its WT value leads to a substantial (~ 2x) decrease of the TAD density in the central nucleus region and, at the same time, to a 20% increase of the chromatin density peak at the NE.

**Figure 7.**
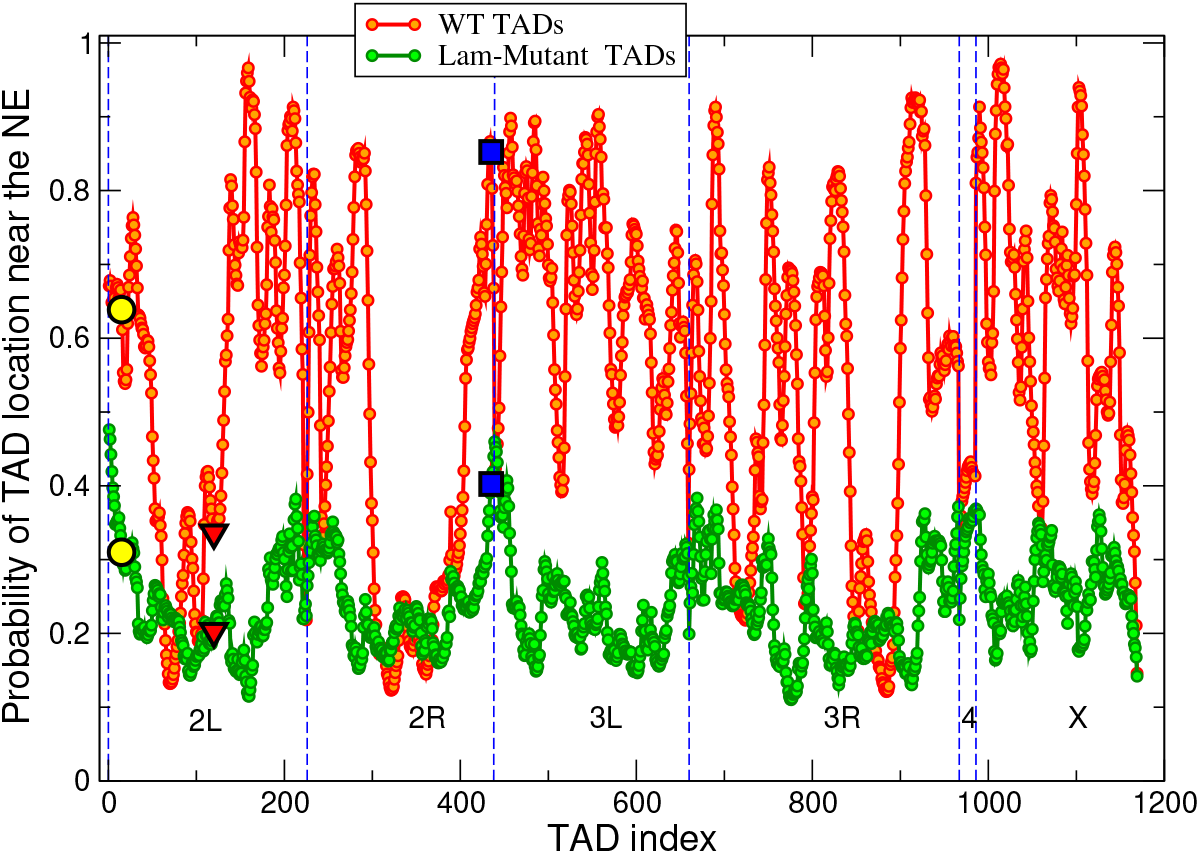
Probabilities of TADs to be found within 0.4 *μ*m layer adjacent to the NE (half the nuclear volume) for the WT (red) and Lamin mutant (green) model nuclei. Statistical error bars are smaller than symbol size. The WT TADs can be partitioned onto two major groups: the TADs with the probabilities greater than 0.5 and the TADs with the probabilities in the range 0.1-0.5. The TADs in the second group belong to the sections of chromosomes containing relatively long continuous stretches (10 – 16 TADs) without LADs. Null L-TADs #15, analyzed in [35] as cytological region 22A, is marked by yellow circles (for the WT and Lamin mutant nuclei). Null L-TADs #120, described in [35] as cytological region 36C, is marked by red triangles. PcG L-TAD #435, analyzed in [35] as cytological region 60D, is marked by blue squares. These three regions have been experimentally shown to detach from the NE in the Lamin mutant nuclei [35]. Our model agrees with the experiment for these three L-TAD regions, providing an additional validation for the model. (Also see model derived cumulative frequencies of radial positions of these regions in Fig.12 in the SI for details).

A more interesting behaviour can be observed when LAD-NE affinity is decreased. The 0.5 *kT* decrease of the affinity results in a substantial reduction of the density peak value near the NE relative to the chromatin density in the central nucleus region (from 1.8 to 1.1) and its 0.2 *μ*m shift from the NE. A further decrease of the affinity to 3 *kT*, which is only a 25% change, leads to a complete transformation of the chromatin density profile to the Lamin mutant-like profile: the density peak at the NE disappears, a substantial amount of the chromatin moves away from the NE, the interior density increases by 80% compared to its WT value. The summary of these effects is presented in Table 1. To the best of our knowledge such a sensitivity of chromatin distribution on the strength of LAD-NE interactions was not observed in the previous studies of mammalian nuclei [54].

The strong dependence of the chromatin density profile on the decrease of LAD-NE affinity (see SI, Fig. 7 (top panel)) suggests that both the number of LADs in the fruit fly genome and their affinity to the NE are likely “tuned” to be rather close to values corresponding to a transition between the Lamin mutant-like and WT-like chromatin distributions.

#### Effects of TAD-TAD interaction variation

We described in “Methods” how we selected the values of the TAD-TAD interaction parameters which produce a good agreement with the experimental Hi-C map [1, 26] and LAD distribution [42]. Here we will discuss what effects the deviations from those optimal parameters will have on the chromatin distribution and the TAD-TAD contact probabilities. The summary of these effects is presented in Table 1.

##### Active-Active TAD interactions

Small increase (from 0.1 to 0.5 *kT*) does not lead to a noticeable change in the TAD radial distribution. Small variations in both directions of intra-arm and inter-arm TAD-TAD contact probabilities are observed.

##### Null-Null TAD interactions

Decreasing the amplitude of these interactions from 1.5 *kT* (WT value) to 0.1 *kT* produces a small effect on the radial chromatin distribution – a 20% increase of the chromatin density in the central nucleus region, and a small decrease of the density near the peak at the nucleus periphery, Fig. 7 in the SI (bottom panel). Similar effect of partial chromatin redistribution toward the nuclear center upon reduction of the attraction between B-type beads has been observed in the single chromosome model of chromatin described in Ref. [51]. This small redistribution of the chromatin in our model results in a minor decrease of intra-arm and inter-arm TAD-TAD contact intensities. On the other hand, the increase of the Null-Null interactions by 1 *kT* (to 2.5 *kT*) leads to a substantial shift of the chromatin from the NE decreasing the fraction of L-TADs at the NE from 25% to 16% by transforming a more narrow main peak near the NE at 1.5–1.9 *μ*m into a wider peak at 0.9 –1.7 *μ*m, Fig. 7 in the SI (bottom panel). Similar effect of the shift of chromatin density toward the central region upon increase of the mutual attraction between B-type beads, when some of them have affinity to the NE, has been observed in the one-chromosome chromatin model described in Ref. [58]. At the same time, the effect predicted in Ref. [51] is the opposite, at least at the same ratio of LAD-NE to B-B attractions as used in our model.

The shift of the chromatin density peak in our model when increasing the Null-Null attraction to 2.5 *kT* (see Fig. 7 in the SI (bottom panel, blue line)) is accompanied by a noticeable increase of intra-arm and inter-arm TAD-TAD contacts probabilities with a reduction of Pearson’s correlation coefficient with the experimental Hi-C map from 0.956 to 0.934. Since 228 of 492 Null TADs contain LADs (the most enriched in LADs class of TADs, see Fig. 1 in the SI), these changes suggest a competition between the LAD-NE and Null-Null (TAD-TAD) interactions for the “optimal” chromatin structure. For comparison, only 54 of 494 Active TADs contain LADs, and the other two less numerous TAD classes contain even smaller numbers of L-TADs (50 of 131 PcG TADs and 18 of 52 HP1 TADs). This suggests that the competition between their TAD-TAD and LAD-NE interactions seems to be less important in the formation of the radial chromatin structure.

##### PcG-PcG TAD interactions

As in the case of Null-Null interactions, the decrease (from 1.5 to 0.1 *kT*) of the attraction between PcG TADs results in minor changes in the chromatin distribution and TAD-TAD contact probabilities. Due to the relative paucity of the PcG TADs we did not pursue further analysis of their selective influence on the chromatin structure.

##### Cross-type TAD-TAD interactions in Lamin mutant vs. WT

Natural separation of TADs into active and inactive compartments suggests that cross-type TAD-TAD interactions would be detrimental to the cell. Does the WT level of LAD-NE affinity make chromatin density profile robust to changes in strength of cross-type TAD-TAD interactions? Decreasing the cross-type interactions from 0.5 *kT* (WT value) to 0.1 *kT* produces practically no change in the WT nuclei and minor changes in the Lamin mutant chromatin distribution (10% – 15% density decrease in the central regions due to small expansion of the chromatin toward the NE), Fig. 6. A small decrease of intra-arm and inter-arm TAD-TAD contacts is observed in both WT and Lamin mutant nuclei. On the other hand, increasing these interactions by 0.5 *kT* (to 1.0 *kT*) results in a substantial (80% – 100%) increase of the chromatin density in the central regions of the Lamin mutant due to further compaction of the chromatin in the globule-like state. The same change of the interactions in the WT nuclei leads to only a small increase of the density (about 20%) in the central regions, Fig. 6. A significant increase of intra-arm TAD-TAD contacts and a moderate increase of inter-arm TAD-TAD contacts are observed in both WT and Lamin mutant nuclei, as it can be seen on the Hi-C map differences (see Fig. 8 in the SI).

The substantial chromatin density changes in the Lamin mutant and the small corresponding changes in the WT nuclei demonstrate that in the absence of the attractive LAD-NE interactions the chromatin distribution is very sensitive to the small changes in the cross-type TAD-TAD interactions. The reduction of this sensitivity in the presence of LAD-NE interactions suggests a stabilizing role of these interactions in maintaining native chromatin distribution in the WT nuclei and preventing cells from potentially negative effects of cross-type TAD-TAD interactions.

##### Role of LAD-NE interactions and highly non-homogeneous LAD distribution along the genome in positioning of TADs relative to NE

To investigate the dynamic positioning of individual TADs within the nucleus relative to the NE, and the role of LAD-NE interactions in this positioning, we have partitioned the nucleus into two spherically symmetric compartments of equal volume: the central spherical region of 1.6 *μ*m radius, and the spherical layer (0.4 *μ*m thickness) adjacent to the NE. In the WT nuclei this 0.4 *μ*m layer contains about 70% of all L-TADs. The probabilities of individual TADs to be in that adjacent to the NE layer for the WT and Lamin mutant nuclei are shown in Fig. 7.

##### Probability of a TAD to be near the NE is determined by a highly variable local linear density of L-TADs (LADs) along the genome

One can see that in the WT nuclei the TADs are partitioned into two major groups: TADs that have high (0.5–0.97) probability to be in the 0.4 *μ*m layer near the NE (outer half the nuclear volume), and the ones that tend to stay away from the NE, with the probability in the range 0.1–0.5 to be in that “near NE” layer. The TADs in the second group are in the sections of the chromosomes containing relatively long continuous stretches (10–16 TADs) without LADs. Quantitatively, one can characterize linear L-TAD density along a chromosome chain (L-TAD frequency of occurrence), *f_L_,* as the ratio of the numbers of L-TADs to the number of all TADs in a given chromosome section.

We see a clear correlation between the probability (*p_NE_*) of a TAD to be in the 0.4 *μ*m “near NE” layer and the *f_L_* value (see Fig. 8). For example, the 1-st dip in the probability distribution for the 2L-arm TADs from #70 to #72 (*p_NE_* = 0.13, Fig. 7 and Fig. 8) is in the stretch of 17 TADs with only a single L-TAD (#78), and has the L-TAD density *f_L_* = 0.05. This density is 6 times lower that the genome-average value *f_L_* = 0.30. On the other hand, the regions in the first group (high *p_NE_*) have a noticeably higher than average L-TAD density: in 2L-arm – TADs from #156 to #166 (*p_NE_* > 0.8) have the average *f_L_* = 0.55.

**Figure 8.**
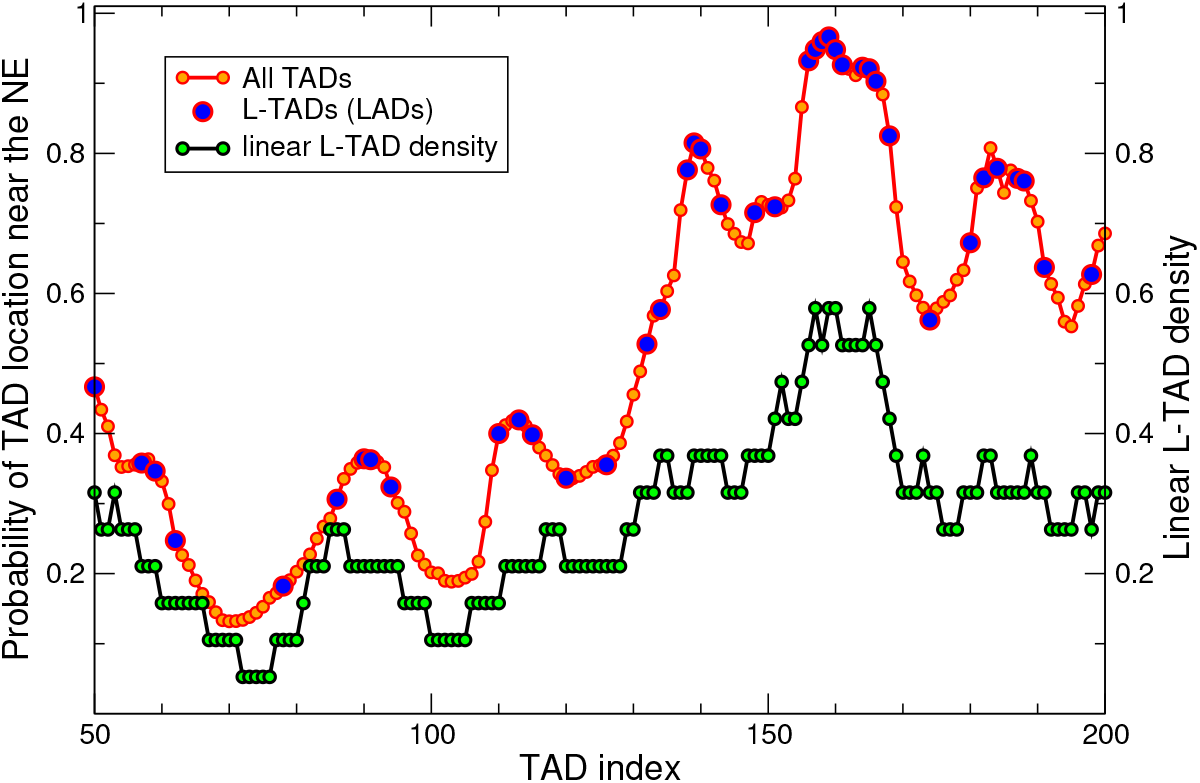
Probability of a TAD to be near NE is largely determined by the linear density of L-TADs surrounding this TAD. Shown are the probabilities of TADs/L-TADs to be found within 0.4 *μ*m layer adjacent to the NE (upper trace), *p_NE_*, and the linear L-TAD density, *f_L_*, along the chromosome chain (lower trace). For visual clarity, only a fraction of 2L chromosome is shown.

This stable separation of the average radial positions of different TADs suggests that the chromatin, despite being in a liquid-like state, has some “averaged global structure” determined by the irregular distribution of LADs (L-TADs) along the chromosomes.

In the Lamin mutant nuclei, the probabilities of TADs to stay closer to the NE (in the 0.4 *μ*m “near NE” layer), presented in Fig. 7, are in the range 0.1–0.4. The overall reduction of these probabilities compared to their WT values, reflecting the change of the TAD’s average radial positions, is in agreement with the detachment of the chromatin from the NE and its compactization in the Lamin mutant nuclei seen on the chromatin density distributions, Fig. 4.

##### Mobility of LADs. The nature of their stochastic distribution

It is well-known that the subset of LADs found at the NE differs substantially from cell to cell. Here we ask if LADs in an individual fruit fly nucleus may also be mobile, and if so, to what extent.

There are 412 LADs in *D. melanogaster* nucleus [46]; they are approximately evenly distributed along the three largest chromosomes: X, 2 and 3. As discussed in “Methods”, we have mapped these LADs onto the 1169 TADs [1] and have found LADs in 350 TADs (L-TADs). The WT value of the LAD-NE affinity (4 *kT*) in our model leads to the 25% fraction of L-TADs being, on average, in the nearest to the NE layer. Increasing or decreasing the LAD-NE affinity relative to its WT value in our simulations leads to an increase or a decrease of this fraction (see Fig. 4 in the SI). This sensitivity (~8% per 0.5 *kT* change) suggests a dynamic balance between the number of L-TADs attached to the NE at any given moment and the rest of L-TADs.

##### LADs in individual nuclei are highly dynamic (mobile)

We find that L-TADs can frequently attach to and detach from the NE during the interphase, and are not permanently anchored to the NE in any given cell. Visualization of motion of five randomly selected L-TADs in each chromosome shows that they are indeed highly mobile on time-scale of 20 minutes (see Fig. 10 in SI and the corresponding movie). One can see that, unlike larger LADs in relatively large human nuclei [48, 47, 34], fruit fly LADs can quickly move though a significant portion of the nuclear volume.

**Figure 9.**
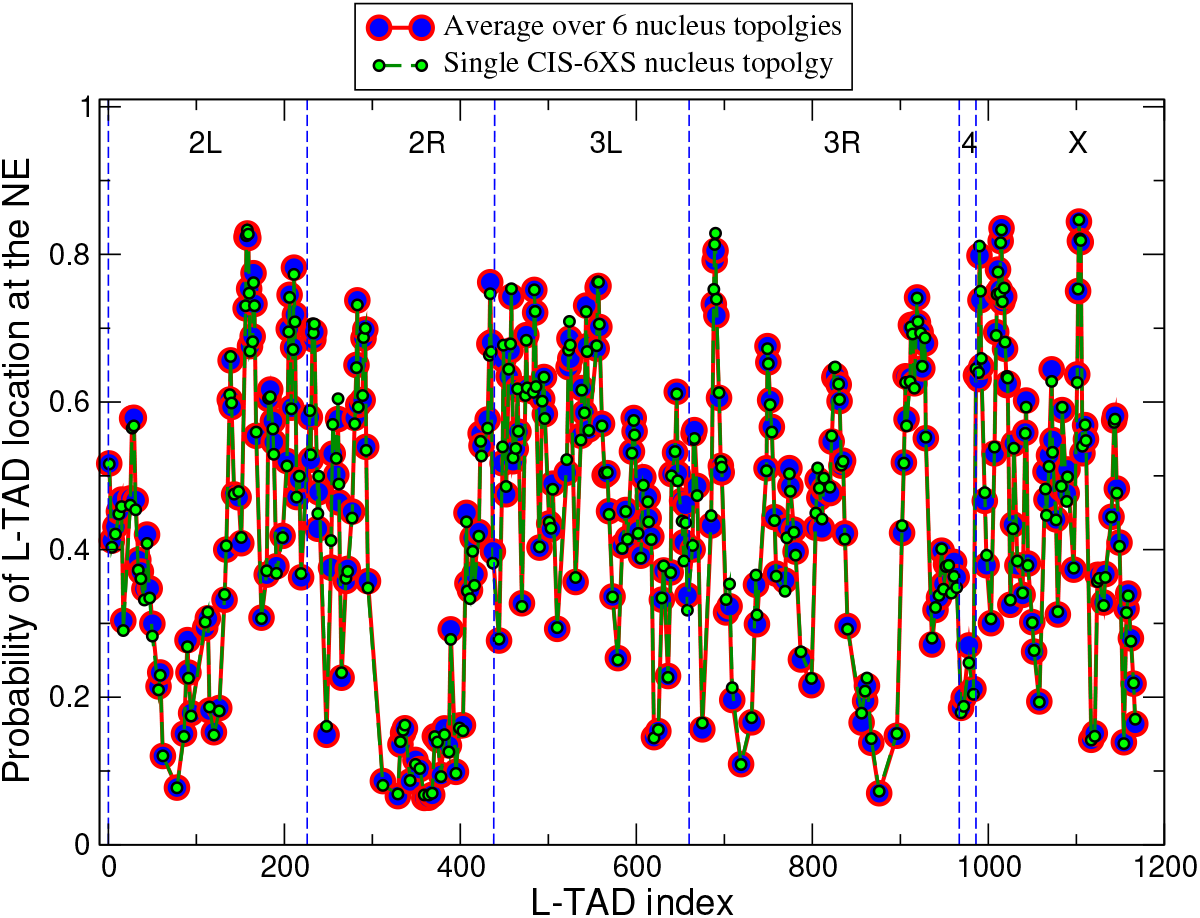
L-TADs are highly mobile within a single cell. Shown are the probabilities of L-TADs to be in contact with the NE. None of the L-TADs stay at the NE all the time, the maximum probability being only 0.85, suggesting a dynamic nature of all LAD-NE contacts. Large red/blue circles are the averages over 6 nuclei with 4 different nucleus topolgies (4 CIS and 2 TRANS). Smaller black/green circles are the probabilities for a single CIS-X6S nucleus topology. All nuclei are of 2 *μ*m radius. Statistical error bars are smaller than symbol size. Despite all 350 L-TADs in the model having the same affinity to the NE, there is a significant spread of their probabilities to be near the NE. A non-negligible number of L-TADs, 12%, have a very low (less than 0.2) probability to be at the NE.

**Figure 10.**
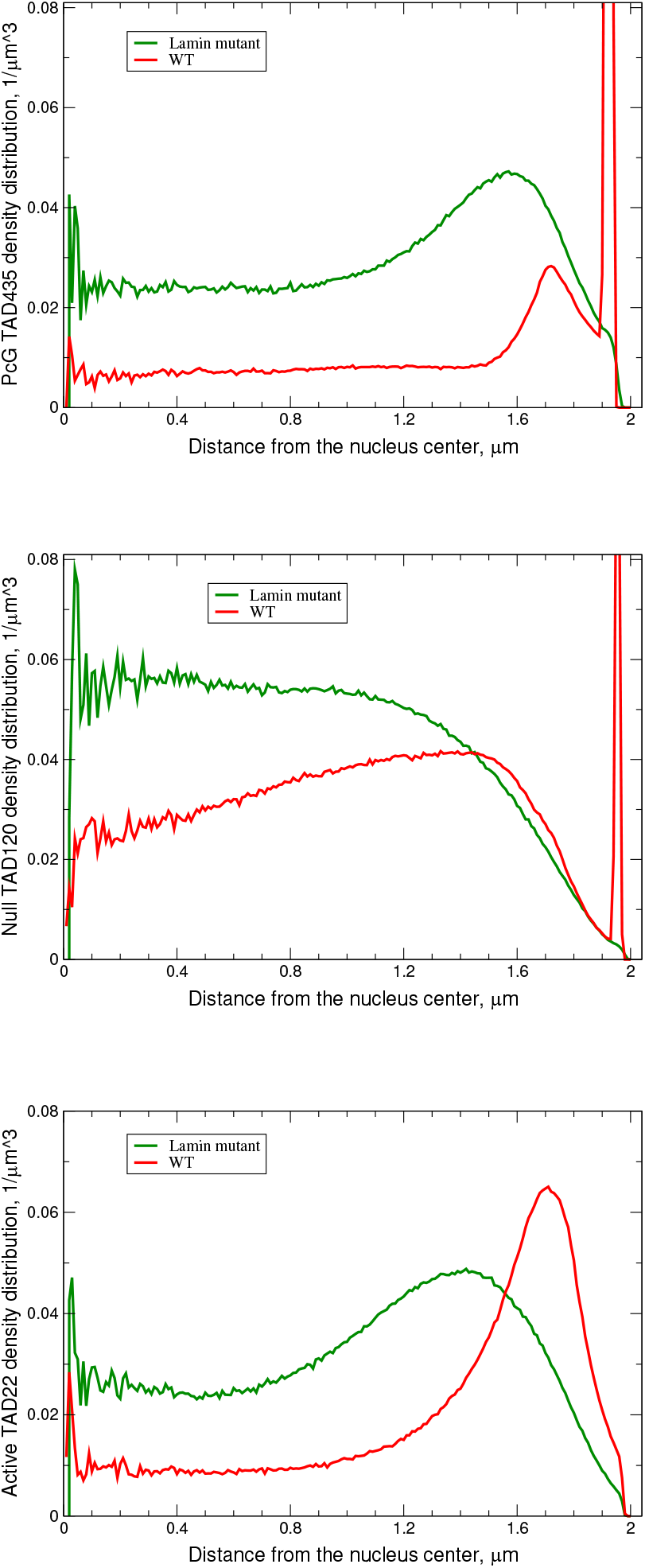
Examples of qualitatively different predicted radial distributions of several TADs, and their responses to lamin depletion. **Top panel:** Radial distribution of the PcG L-TAD #435 (cytological region 60*D* in Ref. [35]) in the WT and Lamin mutant nucleus models. **Middle panel:** Radial distribution of the Null L-TAD #120 (cytological region 36*C* in Ref. [35]) in the WT and Lamin mutant model nuclei. **Bottom panel:** Radial distribution of the Active TAD #22 in the WT and Lamin mutant model nuclei. The distributions of L-TADs (LADs) are bi-modal: the NE-bound mode is characterized by a high and very narrow density peak right at the NE, and the diffusive mode exhibits a noticeable LAD density in the nuclear interior. The bi-modality is consistent with the mobile nature of LADs.

To confirm the suggestion about a highly dynamic nature of LADs on the genomewide level, we have computed the probabilities of all the L-TADs to be in contact with the NE (see the definition in “Methods”), Fig. 9.

To distinguish between cell-to-cell stochasticity and LAD mobility within an individual nucleus, we compare the probabilities computed for the ensemble of 6 nuclei with the probabilities computed for a single nucleus. Note that a hypothetical scenario in which L-TADs were static within each nucleus would yield a qualitatively different – binary – distribution of L-TADs for a single nucleus in Fig. 9 compared to the 6 nuclei average: it would look like L-TADs having only 0 or 1 probability to be in contact with the NE.

From our genome-wide model prediction for all of the L-TADs, the probabilities to be in contact with the NE are lower than 0.85. The fact that the probabilities computed for a single nucleus are noticeably less than 1.0 supports the hypothesis that during the interphase all L-TADs (LADs) bind to and unbind from the NE, within one and the same nucleus (see also Fig. 10 in SI and the corresponding movie). Even those L-TADs with the highest level of NE-binding probability are not permanently attached to the NE.

##### Probabilities of L-TADs to be in contact with the NE vary widely

Another interesting observation is a rather wide spread of the L-TAD contact probabilities seen in Fig. 9. Despite the fact that the model makes a simplification by considering all of the L-TAD-NE affinities to be the same (neglecting the variations in LAD sizes), it predicts very different levels of L-TAD-NE contact probabilities for different L-TADs.

Similar to a more general TAD positioning in a relatively wide layer close to the NE (see Fig. 7), we suggest that this large difference in the L-TAD binding – the contact probabilities vary from 0.07 to 0.84 – is mostly related to a very different linear L-TAD density, *f*L**, (shown in Fig. 8) in the chromosome regions surrounding these L-TADs. For example, the *f*L** varies from 0.08 for L-TAD #78 to 0.7 for L-TAD #1015, with the corresponding probabilities to be found at the NE being 0.08 and 0.84, respectively. Note that both *f*L** values are significantly different from the average linear L-TAD density for all four chromosomes, which is 0.30.

##### Bi-modal radial distribution of L-TADs in WT nuclei

To further explore the predicted dynamic nature of L-TAD binding to the NE, we have computed radial distributions of the positions of two L-TADs (PcG L-TAD #435 and Null L-TAD #120) in the WT and Lamin mutant nuclei, see Fig. 10 (top and middle panels). One can see that for the WT nuclei, where L-TADs have 4 *kT* affinity to the NE, the distributions are bi-modal: they have a bound mode, characterized by a large narrow density peak at the NE, and a diffusive mode with a noticeable density in the nucleus interior. In the case of the Null L-TAD #120 (Fig. 10, middle panel), the WT diffusive mode has a greater amplitude, close to the average single TAD density in the nucleus (~ 0.03 *μ*m^−3^). In the case of the PcG L-TAD #435 (Fig. 10, top panel), most of the WT diffusive mode has a much smaller amplitude (~ 0.01 *μ*m^−3^) with more of the TAD position density localized in the bound mode. These bi-modal radial L-TAD distributions demonstrate further, that in the WT nuclei LADs are not permanently bound to the NE, but are instead rather mobile, being able to attach and detach during the interphase. Similar bi-modal distributions of LADs have been observed in a polymer model of a 81 Mbp genomic region of human chromosome 5 [55].

In addition to L-TADs, the shift of the radial positioning toward the NE in the WT nuclei is also observed for most other TADs (see Fig. 7). A typical behaviour of radial distributions of these TADs is demonstrated using as an example the distribution for the Active TAD #22, depicted in Fig. 10 (bottom panel). The wide distribution peak of this TAD in the Lamin mutant nuclei is shifted toward the NE in the WT nuclei, which is in agreement with different probabilities of this TAD to be within the 0.4 *μ*m layer near the NE (half the nuclear volume) for the Lamin mutant and WT nuclei: 0.30 and 0.62, respectively.

An interesting exception related to this general chromatin density shift toward the NE in the WT nuclei (compared to the Lamin mutant nuclei) are small groups of TADs (e.g., TADs with IDs in the ranges 64–80 and 312–371 in 2L and 2R chromosome arms) that demonstrate small shifts in the radial density distributions toward the NE in the Lamin mutant. We suggest that this opposite pattern is related to a very low linear L-TAD density in these selected regions of chromosomes and, accordingly, a very low averaged attraction to the NE of these regions, compared to the neighbouring regions with higher L-TAD densities. Due to a competition among L-TADs for a limited space near the NE in the WT nuclei, these special groups of TADs with a very low averaged attraction to the NE are effectively pushed away from the NE. Consistent with the above explanation, the differences in the attraction towards the NE are eliminated in the Lamin mutant nuclei, leading to a much smaller spread of TAD probabilities to be near the NE. The remaining small variation – upturns in the green trace at the ends of each chromosome arm in Fig. 7 – is consistent with a greater probability to be at the nuclear periphery for the centromeric and telomeric regions of chromosomes.

The distributions of radial TAD positioning discussed above further suggest that the TADs are highly dynamic in both WT and Lamin mutant interphase nuclei.

## Discussion

This work has two types of outcomes: methodological advances and biological predictions. The latter are focused on the role of interactions between structural components of chromatin such as TADs, as well as their interactions with the NE, in determining the local structure and 3D global architecture of chromatin, and their stability.

### Methodological Advances

We have developed a novel coarse-grained “beads-on-a-string” model of chromatin of the entire *D. melanogaster* interphase nucleus at TAD resolution (~100 kb). One of the major considerations in choosing the TAD level of coarse-graining is that TADs are conserved, stable units of the fruit fly chromatin, interactions between which determine compartmentalization of the chromatin into active euchromatin and more densely packed inactive heterochromatin [1, 15, 106] and, together with the LAD-NE interactions, its global distribution.

The physics-based, as opposed to purely data-driven, approach taken here allows us to answer many “what if” questions hard to address experimentally, focusing on the role of the chromosome–NE interactions on the global and local chromatin structure and its stability, which are the subject of many recent experimental studies [131, 39, 132, 35, 54, 133, 117, 90].

Compared to many existing physics-based models of chromatin in higher eukaryotes, our approach has several methodological novelties; these make a tangible difference with respect to biologically relevant predictions our model can make.

First and foremost, we are simulating an entire biological system – an ensemble of nuclei, corresponding to the experimentally observed set of mutual spatial arrangements of chromosome arm (CIS and TRANS), properly weighted according to experiment. As we have shown, considering the entire biological system *within a physics-based model* proves important to reproducing experiment, specifically to obtain a better agreement with the experimental Hi-C map derived from a very large set of nuclei. A related novel aspect of the developed model is that we consider four distinct epigenetic classes of TADs as four corresponding types of beads, as well as two additional bead types – pericentromeric constitutive heterochromatin (HET) and centromeric regions (CEN) beads.

The second key methodological novelty of our modeling approach is that we are able to simulate temporal evolution of the structure of all chromosomes in the nucleus on the time-scale of the entire interphase. That a computer simulation of an entire fruit fly nucleus at ~100kb resolution can reach the biological timescales of 11 hrs or so is not entirely obvious. Here, we have successfully adapted to the field of chromatin simulations an approach developed in Ref. [120] – implicit solvation with low Langevin friction term. This approach was found successful in the study of protein folding and similar problems in structural biology that faced the same problem: biologically interesting time-scales are far out of reach of traditional simulation techniques often used in this field. While we can not map the resulting time-scales precisely onto real biological time, we are confident that what we call “minutes” in our simulations are not hours or seconds of real biological time (see Fig. 3 in SI).

To arrive at the key interaction parameters that define the behavior of the model, we have used three major criteria, or “rules”: (1) maximizing Pearson’s correlation coefficient between model derived TAD-TAD contact probability map and the experimental Hi-C map [1, 26]; (2) a match between the model derived and experimental set-averaged fractions of LADs contacting the NE [42]; (3) the conditions for the chromatin A/B compartmentalization – the Flory-Huggins phase separation criterion [125]. The criterion (2) is likely a novelty in the development of coarsegrained models of chromatin, which leads to important biological conclusions, see below.

The proposed model has been validated extensively against multiple experimental observations and trends, *independent* from those used in the model construction. A few examples are summarized below. The model derived TAD-TAD contact probability map reproduces all the main qualitative features of the experimental Hi-C maps obtained from *Drosophila* cells [1, 15]. These include increased interactions within chromosomal arms, long-range chromatin contacts visible as bright spots located off the main diagonal, genome compartmentalization manifested as plaid-patterns of TAD-TAD contacts, and the Rabl-like configuration represented by interactions between chromosome arms as “wings” stretched perpendicular to the main diagonal.

Further, the model reproduces the key qualitative results from Ref. [35] that lamin depletion enhances interactions (increases contact frequency) between active and inactive chromatin and leads to a chromatin compaction. Experimentally observed detachment of several cytological regions from the NE in lamin depleted nuclei [35] and their radial positioning are also faithfully reproduced by the model.

In another independent validation of the model, we have demonstrated that it reproduces, *automatically*, experimental chromatin density profiles [117] of both the WT and Lamin mutant nuclei, including some highly nuanced features such the “flatness” of the Lamin mutant density profile away from the NE, Fig. 4, as opposed to, e.g., continued growth of the density toward the center of the nucleus that can be seen when the model parameters deviate from their optimal values, Fig. 6. The fact that the model can reproduce this nuanced behavior is non-trivial, as small variations of the model parameters destroy the agreement (while still predicting the more trivial behavior of the chromatin moving away from the NE in the lamin depleted nuclei).

In summary, the model reproduces about ten distinct features of *Drosophila* interphase chromatin observed in several independent experiments not used in model construction.

### Biological predictions and speculations

Our first noteworthy conclusion is that the positioning of all LADs in *D. melanogaster* interphase nuclei is highly dynamic (mobile) – on the time scale of the interphase the same LAD can attach, detach, move far away from the NE and then re-attach itself to the NE multiple times. This prediction is supported by multiple computational experiments. In particular, the analysis of the distributions of radial positions of single L-TADs: the distributions have two modes - NE-bound and diffusive. Consequently, none of the L-TADs spends all of the time at the NE. This conclusion goes beyond what is known from experiment for fruit fly nuclei: that LADs found at the NE differ from cell to cell. What we show is that, in *any given cell nucleus*, LADs are highly dynamic. We argue that this prediction is robust, as it is an inevitable consequence of the relatively low strength of the LAD-NE attraction. The specific value, 4 *kT*, of this attractive energy used by our model is not arbitrary, it is derived from the experimental fact that only a certain, limited fraction of LADs is on average bound to the NE [42]. A hypothetically much higher value of the LAD-NE affinity that would “glue” all or most LADs to the NE would be inconsistent with the experimental data used to construct the model.

It is worthwhile to compare our genome-wide predictions for LAD mobility in fruit fly with the corresponding experimental findings, which, to the best of our knowledge, are available for human nuclei [34]. In both cases, LADs are not static, but in human nuclei, LAD movements are confined [34] to a relatively narrow layer near the NE, while in fruit fly we see a relatively higher mobility over-all, with many L-TADs traversing the nucleus, from the NE to the center. A more detailed analysis is warranted to quantify the similarities and differences between the nature of LAD mobility in these two organisms.

Observations of the dynamic nature of LADs in interphase nuclei raise a question of the effect of a LAD being in close proximity to the NE on the expression of the genes within that LAD. More specifically, does the expression level vary as the LAD moves between the periphery and the other parts of the nuclear interior? A study systematically tested mammalian promoters moved from their native LAD location to a more neutral chromatin environment and to a wide range of chromatin contexts inside LADs [134]. The study has demonstrated that it is the features encoded in the promoter sequence and variation in local chromatin composition that determine gene expression levels in LADs [134]. If the interplay between promoter sequence and local chromatin features is sufficient to determine the level of transcription inside LADs, then gene expression may be robust to the dynamic nature of LADs, at least in WT nuclei. Future genome-wide studies of the spatial-temporal transcription inside the nucleus may answer this question; combining experiment and computer modeling may be beneficial.

Related to the above conclusion about the dynamic nature of LAD binding to the NE is the prediction that, despite all of the L-TADs in our model having exactly the same affinity to the NE, the probability of L-TADs binding to the NE varies by an order of magnitude between L-TADs. We explain this variation by the corresponding pronounced variation, up to 9 times, of the local linear L-TAD density along the chromatin chains, in contrast to an earlier suggestion that it is the highly variable LAD-NE affinities of relatively large LADs in human cells that may be responsible for the differences in the frequency of LAD binding to the NE [50]. A potentially biologically relevant consequence of our finding is that the genetic/epigenetic features of a given TAD alone can not fully determine its fate with respect to probability of being found near the NE, even if the stochastic component of the positioning is eliminated by averaging over time and an ensemble of nuclei. The distribution of LADs along the genome strongly affects the average radial positioning of individual TADs, playing a notable role in maintaining a non-random average global structure of chromatin, within its over-all liquid-like state.

We also find that the specific strength of the WT value of LAD-NE attraction puts the chromatin very near the “phase boundary”, separating two qualitatively different chromatin density distributions; a mere 12% (0.5 *kT*) decrease of the LAD-NE affinity strength changes the shape of the chromatin density profile appreciably, from the WT one to one that resembles the Lamin mutant density profile. Changing the LAD-NE affinity by 25% from its WT value (1 *kT* decrease) results in a drastic (60%) decrease in the fraction of L-TADs at the NE. One proposed biological consequence of being on the “phase boundary” is as follows. If we assume that the ~ 12% (0.5 *kT*) variation in LAD-NE affinity occurs naturally, then the high sensitivity of the chromatin structure to the strength of LAD-NE affinity might explain variability of chromatin architecture between nuclei of the same tissue and between different tissues. Indeed, some LADs are conserved between cell types, while others are more variable [135]. LADs that display less consistency between cells in a population tend to be specific to cells where genes, located in these LADs, are transcriptionally repressed [136]. Cell type-specific genes located in variable LADs are released from the NE upon cell type differentiation [48].

Another set of model predictions focuses on the potential role of LAD-NE interactions in the sensitivity, or lack thereof, of chromatin 3D architecture to other key interactions (TAD-TAD), which together create the delicate balance that determines the nuclear architecture. Recent studies [26, 137, 82, 54, 51], including this one, leave little room to debate the importance of LAD-NE interactions in genome organization. As in previous works, *e.g.* on mouse [54], agreement of the polymer model with experiment can only be achieved in a rather narrow window of parameters that determine TAD-TAD and LAD-NE interactions in fruit fly nucleus. Our model goes further, by allows us to differentiate between the four main types of TADs: we find that among transcriptionally repressed TAD types, Null-Null interactions have the strongest effect on the 3D chromatin architecture. Also, as previously reported, one must assume a relatively weak mutual attraction between Active-type TADs. The fact that very different models applied to very different organisms, from fruit fly to mammals [54, 51], arrive at several similar general conclusions regarding the role of the interplay of the interactions between chromatin units and the NE, speaks for a certain degree of conservation of chromatin organization across species. We would like to note, however, that in a resent computational study of mammalian nuclei [54], a much larger (compared to our work) relative variation of model LAD-NE affinities (up to 3 times) was found compatible with the chromatin distribution in the WT nuclei.

In contrast to the predicted effect of small changes in LAD-NE interactions on the radial chromatin distribution, this distribution is rather insensitive to even relatively large (30–100%) changes in the strength of the interactions between TADs. The changes in the interactions between the Null TADs have the most effect, but even that effect is significantly smaller than the changes in chromatin density profile resulting from similar relative changes in the LAD-NE affinity.

Critically, in contrast to WT nuclei, chromatin density distribution in the Lamin mutant is predicted to be sensitive to increase of the strength of cross-type TAD-TAD attractive interactions. This comparison suggests that another role of relatively strong LAD-NE interactions is in providing a stable global environment with a low sensitivity to small fluctuations in TAD-TAD interactions. We speculate that the low sensitivity of global chromatin architecture (radial chromatin density distribution) to TAD-TAD interactions hints at the possibility of a mechanism in which changes in TAD-TAD interactions can be more important for local regulation of gene transcription activity. We also make a testable prediction that many aspects of the chromatin architecture will be more variable among cells with fewer LAD-NE contacts and even more so in the Lamin-depleted cells. These may include TAD-TAD contacts, chromatin radial distribution, Rabl configuration, spatial segregation of chromosome territories. In fact, a greater variability of the chromatin radial distribution in proventriculus nuclei of the *Drosophila* Lamin mutants in comparison to WT nuclei can be observed by comparing three experimental groups ([117]). Thus, a dramatic loss or dysfunction of lamins during aging or disease [48, 138, 61, 62, 63] may contribute to the increased disorder in gene expression due to greater variability of the global chromatin architecture predicted in this work.

Turning off LAD-NE interactions (simulating Lamin mutant) results in almost complete detachment of the chromatin from the NE and its compaction, causing a 2-fold increase of the chromatin density in the central region of the nucleus. At the same time, complete removal of the NE in the model results in chromatin decompaction and separation of the chromosome territories on very short time-scale, orders of magnitude shorter that the duration of the interphase. This result suggests that, without the NE, the interactions between TADs are not strong enough to keep the chromatin of the Lamin mutant in a globule-like form on biologically relevant time-scales of hours. At the onset of mitosis, the NE is disassembled and the entire genome is condensed into mitotic chromosomes [139]. In organisms with an open mitosis, such as *Drosophila* (except syncytial embryonic divisions), NE reformation occurs by recruitment of nuclear pore complexes and membrane components to the surface of the segregating chromosomes. The process begins in late anaphase with the binding of nuclear pore complex proteins to chromosomes and is completed with the recruitment and fusion of membranes during telophase [140]. Thus, the enclosing role of the NE is already established at the beginning of the interphase. This process could have evolved to prevent chromosomes from further unfolding and detaching from each other.

Taken together, these observations suggest a dual mechanical role of the NE in the WT nuclei: it is not simply a confinement or “stretcher” of the chromatin, but, rather, the NE acts as an “attractive enclosure”, which simultaneously expands and confines the chromatin, and stabilizes both its local and global 3D structure.

Contrary to the chromatin density profiles, which reflect the global 3D chromatin architecture, the predicted Hi-C maps are not very sensitive to the changes in the LAD-NE affinity, suggesting that local chromatin structure is determined mostly by the TAD-TAD interactions. Indeed, increasing the interactions strength between the most numerous strongly interacting Null TADs by 1 *kT*, from their 1.5 *kT* optimum WT value, leads to a substantial increase in the number of intra-arm and inter-arm contacts. A similar effect on Hi-C maps is predicted for the change in interactions between TADs of different types.

By and large, the predicted Lamin mutant Hi-C map looks rather similar to that of the WT one (Pearson’s correlation coefficient between these maps is 0.9989); this similarity, within each chromosome, was previously observed in lamin knockdown nuclei [35]. A more detailed analysis shows that most TAD-TAD contacts, including inter-chromosome contacts, are slightly enhanced in the Lamin mutant compared to the WT nucleus (see the difference map in the SI, Fig. 8), consistent with the over-all compaction of chromatin observed experimentally in LamA25 mutant [117]. Small areas where the contact frequency decreases in the Lamin mutant are predicted to be limited to close to diagonal intra-arm contacts. These findings are consistent with predictions made for polytene chromosomes in fruit fly [76].

The over-all increase of TAD-TAD contacts and compaction of the chromatin in the Lamin mutant may bring TADs of different chromosomes closer to each other, facilitating inter-chromosome interactions, which is exactly what we observe in our simulations. In particular, the Lamin knockdown increases the chromatin density in a fraction of TADs enriched in active chromatin, and enhances interactions between active and inactive chromatin [35]. Using our model, we can go further and quantify some of these changes. From the model Hi-C maps, we have estimated the sums of contact probabilities of each Null TAD (inactive B-type TADs) with the Active TADs in the Lamin mutant and WT model nuclei, see Fig. 11 in SI. We predict a noticeable, 22% on average, increase of these active-inactive chromatin contacts in the Lamin depleted nuclei. The reduction of active-inactive chromatin contacts in WT suggests a stabilizing role of the LAD-NE interactions in maintaining native chromatin distribution and preventing cells from potentially detrimental effects of cross-type TAD-TAD interactions.

Taken together, our modeling data indicate that LAD-NE interactions play a diverse and prominent role in 3D genome organization.

## Conclusions

We have developed a coarse-grained model of *D. melanogaster* interphase nuclei at TAD (~100 kb) resolution that describes dynamics and time evolution of all fruit fly chromosomes, and their interactions with the nuclear envelope (NE), on the time-scale of the entire interphase. The model takes into account different types of TAD-TAD interactions between different epigenetic classes of TADs, attractive interaction between LADs and the NE, and is tuned to reproduce the experimental Hi-C map and the fraction of LADs positioned at the NE. Several methodological novelties proved important to achieve good agreement with experiment, including explicitly accounting for different experimentally observed mutual spatial arrangements of the chromosome arms (nucleus topologies). The model has been validated against multiple distinct features of *Drosophila* interphase chromatin, not used in the fitting of its parameters.

We have used the model to explore, in detail, how several key characteristics of the chromatin 3D architecture, including the overall chromatin density distribution and Hi-C maps, are sensitive to the interaction strength between different classes of TADs, and between LADs and the NE. Some of our general conclusions agree with previous findings based on models of mammalian nuclei, which supports conservation of several general principles of chromatin organization across species.

Multiple genome-wide predictions have been made in this work. We predict a very dynamic nature of binding of LADs to the NE in *D. melanogaster* interphase nuclei. We also predict an increased sensitivity of global chromatin architecture to the fluctuations in TAD-TAD interactions in lamin depleted nuclei compared to the WT, where relatively strong LAD-NE interactions suppress this sensitivity. The proposed model predicts radial positioning of individual TADs in the nuclei, in particular, the probabilities of TADs to be in the high density layer at the NE. We predict a positive correlation between these probabilities and local linear (along the chromatin chain) densities of LADs around TADs, suggesting a significant role of LAD distribution in average 3D positioning of individual TADs.

We conjecture that one important role of the distribution of LADs along the chromosome chains and their attractive interactions with the NE is to create a non-random average global structure of chromatin and to protect its integrity and stability against inevitable cell-to-cell variations in TAD-TAD interactions. We also predict greater variability of the chromatin architecture due to loss or dysfunction of lamins, which may contribute to the increased disorder in gene expression during aging or disease.

## Abbreviations

3D: three-dimensional
CEN: centromeric
Chr: chromosome
HET: pericentromeric constitutive heterochromatin
LAD: lamina-associated domain
L-TAD: TAD that contains LADs
LJ: Lennard-Jones
MSD: mean squared displacement
NE: nuclear envelope
TAD: topologically associating domain
WT: wild-type.

## Acknowledgements

We thank Raju Nadimpalli for help with ESPResSO input scripts, chromatin visualization and Hi-C map processing; Samira Mali for technical assistance and for producing the movie clip of the chromosomes dynamics; Yoonjin Kim for technical assistance; Frank Alber for a useful discussion and for providing a comprehensive set of configurational snapshots from Ref. [26].

## Declarations

### Authors’ contributions

IT and AO designed the dynamical model with input from IS and NK. IT implemented the model with early input from NK. IT, AO and IS analyzed the results. IT, AO, IS and NK wrote the original manuscript.

### Availability of software

The modeling code ESPResSo 3.3.1 [118] used in this research is available at http://espressomd.org/. The software is free, open-source, published under the GNU General Public License (GPL3). Visualization of motion of randomly selected L-TADs in each chromosome on a time-scale of 20 minute is available at https://drive.google.com/drive/folders/1Dwfe7hbCjARxnvhf7oP76sVaTRa0RE6Z

### Ethics approval and consent to participate

Not applicable.

### Consent for publication

All authors read and approved the manuscript.

