## Supplemental Information for "Strong interactions between highly dynamic lamina-associated domains and the nuclear envelope stabilize the 3D architecture of Drosophila interphase chromatin"

NEW RESULTS

### Methods

#### Bead size and mass

The radii value used in this work are modified TAD radii from [1] (scaled by a factor 1.254031 to reflect the double volume (two homologous TADs) of our beads). The radii values estimated in [1] are based on the DNA density in TADs, $\sim$60 Mb/$\mu$m$^3$, and correspond to 12% volume fraction of all euchromatin domains with respect to the nucleus volume [2]. The radii of the HET and CEN domains in [1] are based on the DNA density $\sim$90 Mb/$\mu$m$^3$ and correspond to 3.7% volume fraction [2].

After the re-scaling, the average radius of the 1169 beads representing homologous pairs of TADs in our model is 0.09 $\mu$m.

It was suggested in [3] that the conformational state of chromatin across organisms depends on the chromosome to nucleus volume ratio, which emphasizes the need to make sure that computational models get this critical parameter right, in order to faithfully describe reality.

#### Bead-bead interactions and bead types

Bonded interactions between neighbouring beads along the chromatin chains are modeled by a harmonic potential,

$$U_b(r_{ij}) = \frac{1}{2}\frac{\kappa}{l_{ij}^2}(r_{ij} - l_{ij})^2\,, \qquad (1)$$

where $r_{ij}$ is the center-to-center distance between beads $i$ and $j = i + 1$, the equilibrium bond length, $l_{ij} = r_i + r_j$, is a sum of beads' radii, and $\kappa$ is a bond strength parameter. Following Ref. [4], we choose $\kappa = 10\ kT$ to suppress chain crossing to a large degree. Non-bonded interactions between beads are described by a short-range Lennard-Jones-cosine (LJ-cos) potential [5], with a minimum at $r_{ij} = l_{ij}$ and

$\sigma_{ij} = l_{ij}/2^{\frac{1}{6}}$.

$$U_{nb}(r_{ij}) = 4\epsilon\left[\left(\frac{\sigma_{ij}}{r_{ij}}\right)^{12} - \left(\frac{\sigma_{ij}}{r_{ij}}\right)^{6}\right], \; 0 < r_{ij} \le l_{ij} \quad (2)$$

$$U_{nb}(r_{ij}) = \frac{1}{2}\epsilon\left[cos(\alpha r_{ij}^2 + \beta) - 1\right], \; l_{ij} < r_{ij} \le 3^{\frac{1}{6}} l_{ij},$$

$$U_{nb}(r_{ij}) = 0, \; r_{ij} > 3^{\frac{1}{6}} l_{ij}.$$

At $r_{ij} > l_{ij}$ the LJ-cos function smoothly goes to 0 at $3^{\frac{1}{6}} l_{ij}$, which is fulfilled by the corresponding choice of the parameters $\alpha$ and $\beta$ (see Table 2 in Ref. [4]).

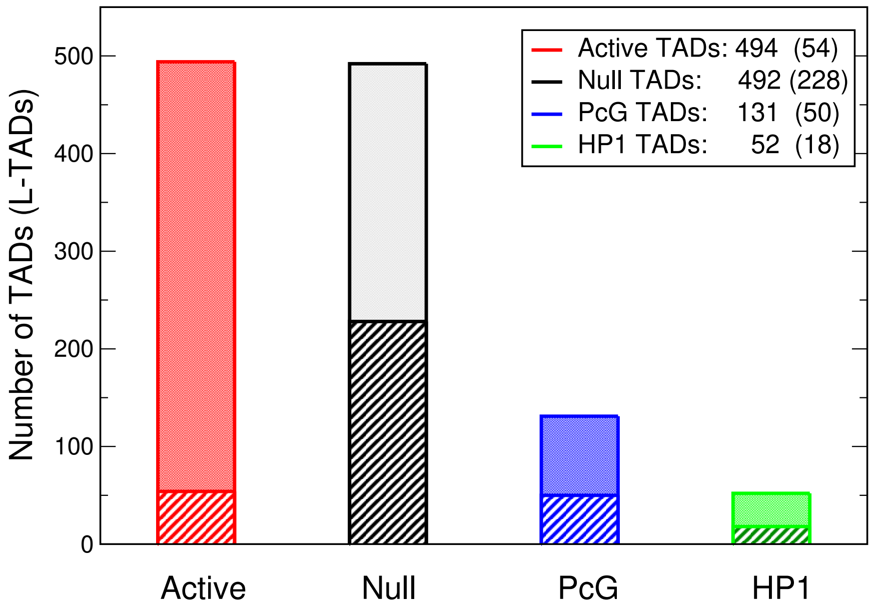


**Figure 1** The number of TADs and LADs containing TADs (L-TADs) in each of the four epigenetic classes of TADs (Active, Null, PcG and HP1) are shown: the numbers of TADs – as solid color bars, the numbers of L-TADs – as bars with diagonal stripes (and numbers in the brackets in the legends).

Following our TAD–bead equivalence, we specify four corresponding bead types: Active, Null, PcG, and HP1. The number of TADs in each epigenetic class and the corresponding number of beads in each bead type in our model are shown in Fig. 1. Non-bonded attractive interactions (Eq. 2) between beads of the same type are characterised by its own, bead-type specific interaction well depth parameter. Attractive interactions (Eq. 2) between the TADs of different types – cross-type interactions – are not type-specific and are characterised by a single well depth parameter.

Repulsive interactions of the nucleolus with all beads, except HET beads, are described by a pure repulsive LJ potential function [5]:

$$U_r(r_{ij}) = 4\epsilon_r \left[ \left(\frac{\sigma_{ij}}{r_{ij}}\right)^{12} - \left(\frac{\sigma_{ij}}{r_{ij}}\right)^{6} + \frac{1}{4} \right], \; 0 < r_{ij} \leq l_{ij}\,, \qquad (3)$$
$$U_r(r_{ij}) = 0\,, r_{ij} > l_{ij}\,.$$

Here $l_{ij} = r_i + r_{nuc}$ is a contact distance between bead $i$ and the nucleolus ($j$) of radius $r_{nuc} = 0.333$ $\mu$m [1], $\epsilon_r = 3$ $kT$, and $\sigma_{ij} = l_{ij}/2^{\frac{1}{6}}$.

To localize HET beads near the nucleolus (see Ref. [1]), their interactions with the nucleolus are set to be strongly attractive, described by a standard LJ potential with $\epsilon_{i,nuc} = 6$ $kT$ well depth and $\sigma_{i,nuc} = (r_i + r_{nuc})/2^{\frac{1}{6}}$.

#### Interactions in specific pairs of remote loci

Experimentally, 268 pairs of remote chromatin loci, with higher than expected contact probabilities, have been identified from the Hi-C data [6]. We map these specific loci pairs onto the pairs of TADs and introduce special interactions between the corresponding beads. The pairs of are ordered according to their degrees of contact enrichment: a ratio of the observed to expected contact probabilities, $P_{ij}^{obs}/P_{ij}^{exp}$. Since it is reasonable to assume that the bead-bead contact probability $P_{ij} \sim exp(-\Delta U_{ij}/kT)$, where $\Delta U_{ij}$ is a well depth of the effective interaction potential between beads $i$ and $j$ in the specific pair, one can express $\ln(P_{ij}^{obs}/P_{ij}^{exp})$ as

$$\ln\left(\frac{P_{ij}^{obs}}{P_{ij}^{exp}}\right) \approx -\frac{\Delta U_{ij}}{kT} + \frac{\Delta U^{std}}{kT}. \qquad (4)$$

Here, $\Delta U^{std}$ is some unknown average "standard" attractive interaction between beads (TADs), assumed in the expected $P_{ij}^{exp}$.

To estimate $\Delta U_{ij}$ we fitted the experimental decay curve (see Fig. 2) of the logarithm of ordered (index $n$) degrees of enrichment for the 268 specific pairs of beads (TADs) with the formula:

$$\ln\left(\frac{P_n^{obs}}{P_n^{exp}}\right) = -1.18(n/267)^{0.2} + C\,, \qquad (5)$$

where index $n$ of the ordered pairs runs from 0 to 267 and parameter $C = 2.22$.

Eqs. 4 and 5 allow us to determine the well depth $\epsilon_n$ of the LJ potential used to describe attractive interactions in each specific pair (in $kT$ units):

$$\epsilon_n = -1.18(n/267)^{0.2} + E_{sp}\,. \qquad (6)$$

Parameter $E_{sp}$ absorbs the constant $C$ from Eq. 5 and the unknown interaction $\Delta U^{std}$ from Eq. 4: $E_{sp} = C - \Delta U^{std}$. We vary $E_{sp}$ between 3 and 6 $kT$ and find that the optimum for the model TAD-TAD contact probability matrix (Hi-C map)

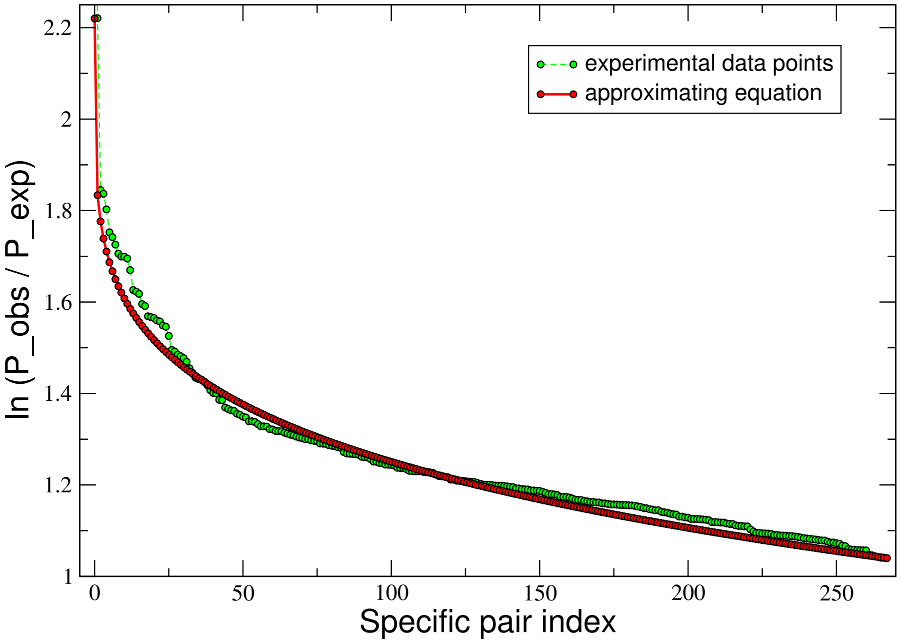


**Figure 2** Determining the interaction strength between the special pairs of TADs. Shown is the logarithm of the ordered (index $n$) degree of enrichment ($\ln(P_n^{obs}/P_n^{exp})$) for the contact probabilities of 268 specific pairs of remote chromatin loci [6] and approximating fit (Eq. 5) used to determine the attractive interaction LJ well depth for the interactions between the beads in that specific pairs.

corresponds to $E_{sp} = 3.5\ kT$. With this value of $E_{sp}$, the attractive interaction well depth $\epsilon_n$ varies from 2.3 to 3.0 $kT$ for most (263 of 268) specific bead pairs.

#### Nuclear envelope and its interactions with chromatin chains

We model the NE as a spherical boundary which confines the motion of the chromosomes and attracts LADs [7, 8, 9, 10]. Three radii of the NE are used: 1.94 $\mu$m, 2.0 $\mu$m and 2.06 $\mu$m. These account for natural cell-to-cell variability; without it, chromatin density profiles near the NE are unrealistic compared to experiment. Following [4], we map positions of 412 LADs [8] (a median size of LADs is about 90 kb) onto the chains of 1169 TADs. If a TAD contains LAD, then the corresponding bead can attractively interact with the NE. After mapping, we have determined 350 TADs (L-TADs) that contain LADs. The fraction of L-TADs with two or more LADs is about 13%. The distribution of TADs among epigenetic classes and the numbers of L-TADs in each class in our model are shown in Fig. 1.

We describe all L-TAD–NE attractive interactions by the LJ-cos potential (Eq. 2) with a single well depth parameter $\epsilon_L$ (neglecting the variations in the LAD-NE affinity due to different LAD sizes). The interaction distance $r_{ij}$ is defined as a distance $r_{in}$ from the bead $i$ center to a spherical surface positioned at $r_n = 0.04$ $\mu$m outside the NE boundary. The energy minimum is placed at the distance $r_i$ (bead $i$ radius) from the NE, resulting in parameters $l_{in} = r_i + r_n$ and $\sigma_{in} = l_{in} 2^{\frac{1}{6}}$.

Fraction of LADs at the NE, which depends on the affinity $\epsilon_L$, is calculated as the average (over the described above ensemble of 18 nuclei) fraction of L-TAD beads within (position of the bead centers) 0.09 $\mu$m layer (average bead radius) at the NE. The thickness of this layer corresponds to about 0.2 $\mu$m layer of beads which are in contact with the NE.

Interactions of the NE with beads not containing LADs are described by the purely repulsive potential $U_r$ (Eq. 3) with $\epsilon_r = 3\ kT$, and with $r_{ij}$, $\sigma_{ij}$ and $l_{ij}$ replaced by the L-TAD–NE interaction parameter $r_{in}$, $\sigma_{in}$ and $l_{in}$, defined above.

#### Lamin Mutant model

The Lamin mutant nucleus model has the L-TAD–NE affinity parameter $\epsilon_L$ reduced to 0.1 $kT$ level. The small but non-zero value is chosen to use the same LJ-cos function (Eq. 2) as for the other values of the L-TAD–NE affinity, and to reflect possible other tethering mechanisms of LADs to the NE [11] in the Lamin B (Dm0) knockdowns or mutants with much smaller levels of LAD-NE attraction than in the WT nuclei. We use Lamin B mutants as the experimental counterpart to our Lamin mutant nucleus model. All other interaction parameters are unchanged from the wild-type (WT) model.

#### Experimental chromatin density profiles

The results of the simulations are compared with the experimental data published previously in Ref. [12]. In that experiment, *D. melanogaster* line with the Lamin B (Dm0) mutant genotype $\mathrm{Lam}^{A25}\mathrm{pr}^1$/CyO; $\mathrm{st}^1$ (BSC, #25092) was used to investigate the role of the NE in nuclear architecture. Proventriculi (the end parts of the larval foreguts) were dissected from the $\mathrm{Lam}^{A25}$ mutant and Canton-S wild-type (WT) 3rd instar larvae. The radial distributions of DAPI-stained chromatin inside the nuclei were measured by quantification of the fluorescence intensity along the nuclei radii using confocal microscopy. The radial coordinates for each nucleus in pixels were translated into relative coordinates and normalized, see Ref. [12]. Mutation $\mathrm{Lam}^{A25}$ removes the CaaX box from Lamin B [13] disrupting the interaction of Lamin B with the NE. As a result, tethering of the chromatin to the nuclear periphery is impaired in the proventriculus nuclei of $\mathrm{Lam}^{A25}$ mutants as seen from the change in the chromatin radial distribution [12].

#### Matching simulation time to biological time

To relate the simulation timescale with the experiment [14], for the chromatin dynamics in *D. melanogaster* interphase nuclei, we compare [4] a diffusive motion of model beads in the simulations with the experimental interphase chromatin diffusion, observed in experiment [14], and use the match to estimate the scaling factor $\lambda$ that converts the simulation time to real biological time. To characterize the diffusion, we calculate time dependencies of the distance $R_i(t)$ between a selected bead $i$ and the nucleus center. Trajectories of for 9 randomly selected beads, which do not contain LADs and, thus, cannot be directly affected by the attraction to the NE, have been extracted. The dependence of the mean squared displacement (MSD) on the time interval $\Delta t$, $\langle \Delta R_i^2(\Delta t)\rangle = \langle [R_i(t+\Delta t) - R_i(t)]^2\rangle$, is calculated for 3 different values of the friction parameter $\gamma$ ($0.01/\tau$, $0.1/\tau$ and $1/\tau$). The averaging is performed over 9 selected beads along 18 trajectories for each value of $\gamma$. We fit the first $300 \cdot 10^3$ time steps of each of the three curves $\langle \Delta R_i^2(\Delta t) \rangle$ (see Fig. 3) with the equation for sub-diffusive motion of chromosomal loci [14], which describes MSD as a function of time interval $\Delta t$,

$$\langle \Delta R_i^2(\Delta t) \rangle = 4D_{app} \left( \Delta t / \lambda \right)^{0.39} , \quad (7)$$

where $D_{app}$ is the apparent diffusion coefficient.

This equation (with $\lambda = 1$ and $\Delta t$ in seconds) describes the experimentally observed diffusion of chromosomal loci over the time periods ranging from 1 to $10^3$ s [14]. We obtain reasonable fits of the simulation derived MSDs with $4D_{app} = 0.061$ $\mu$m$^2$ and $\lambda = 10^4$ for $\gamma = 0.01/\tau$, $\lambda = 2.8 \cdot 10^4$ for $\gamma = 0.1/\tau$, and $\lambda = 20 \cdot 10^4$ for $\gamma = 1/\tau$ (for $\Delta t$ measured in a number of the simulation time steps). These values of $\lambda$ give us the numbers of simulation time steps that corresponds to 1 sec of a real biological time. The scaling translates the length of the $40 \times 10^6$ time step simulations with $\gamma = 1/\tau$ to 3 min of real nucleus time, and long $400 \times 10^6$ time step simulations of WT and Lamin mutant nuclei with $\gamma = 0.01/\tau$ as corresponding to 667 min (11 hrs) of real time.

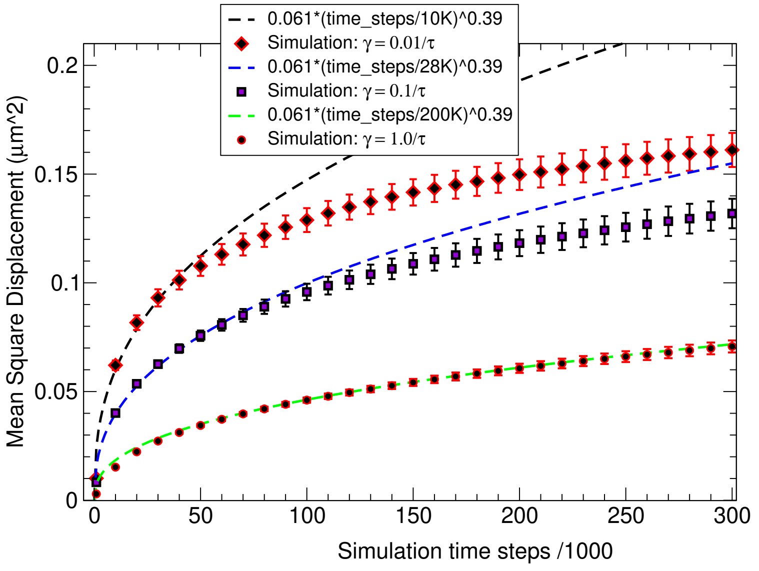


**Figure 3** Time dependence of mean squared displacement (MSD) of bead averaged over 9 beads and 18 trajectories of the model nuclei with different topologies and size, simulated with 3 different values of the friction parameter $\gamma$ ($0.01/\tau$, $0.1/\tau$ and $1/\tau$). The simulation data points (symbols with error bars) are fitted with Eq. 7 (dashed lines) – the equation corresponds to the experimentally observed time dependence of MSD for a chromosomal loci [14]. The fit is used to estimate the simulation time scaling factor $\lambda$, which determines the mapping between the simulation time and biological time of a nucleus. Error bars in each curve are RMSD of 9-bead-average values in 18 trajectories.

### Contact probability (Hi-C) map

The TAD-TAD contact probability map (Hi-C map) of the model is defined as the average of the 18 Hi-C maps (1169x1169) calculated over the 18 trajectories of the model nuclei of different sizes and mutual arrangements of the chromosomes. Each matrix element characterizes a contact probability $p_{ij}$ of the two beads defined as the fraction of the configurations where the beads $i$ and $j$ are in a contact. We define the bead-bead contact as the configuration with the distance between the centers of the two beads $d_{ij} < r_i + r_j + 0.2$ $\mu$m.

### Model development

#### Interaction parameters of the models

Three stages of model development are used. In the initial stage, only HP1 type beads – classical heterochromatin TADs – were included into B-type set of beads. The rest of the beads were characterised by a weak 0.1 $kT$ TAD-TAD attractive interaction. We analyzed 144 combinations of the interaction parameters by generating $40 \times 10^6$ time step MD trajectories for each parameter set, using only one nucleus topology. We scanned the following range of parameters with in 1 $kT$ increments. The attractive LJ-cos interactions: 3 – 6 $kT$ for the $E_{sp}$ parameter in the specific TAD pair interactions; 1 – 6 $kT$ for the HP1-HP1 (B-type beads) interactions; 2 – 6 $kT$ for the LAD-NE (L-TAD–NE) interaction parameter $\epsilon_L$. For each parameter set the Hi-C map was calculated and compared with the experimental Hi-C map [6, 1] resulting in the range of Pearson's correlation coefficients from 0.895 to 0.950. This analysis allowed us to narrow down the ranges of the parameters for further refinement.

In the next stage of our model development, in addition to the HP1 beads, we considered transcriptionally inactive [6] Null and PcG beads as members of the inactive chromatin B-compartments. The corresponding attractive interactions (Eq. 2) between beads of the same type in these B-compartments are characterized by the LJ-cos well depth parameters $\epsilon_B$. To allow possible compartmentalization within each of these three bead types, we describe the cross-type interactions between these beads (Eq. 2) by the same small LJ-cos parameter as the one, $\epsilon_{AB}$, used for the A-B cross-type interactions. Active-Active interactions (Eq. 2) between beads in A-type compartments are described by the LJ-cos parameter $\epsilon_A$. Interactions between HET and CEN beads are considered as A-B cross-type interactions.

Results from the previous stage allowed us to focus on a more narrow range of model parameters: only 47 sets of the interaction parameters. The selected ranges for the LJ-cos interaction well depths are: 1 – 2 $kT$ for $\epsilon_B$; 3 – 4 $kT$ for $E_{sp}$; and 2 – 6 $kT$ for the $\epsilon_L$ (L-TAD–NE interactions). Parameters $\epsilon_A$ and $\epsilon_{AB}$ were fixed at 0.1 $kT$ (to satisfy selection rule #3 above for $\epsilon_{AB}$). For each set, a $40 \times 10^6$ time-step trajectory was generated for a single CIS-X6S nucleus topology (see Fig. 1 in "Methods", main text), the Hi-C map was calculated and compared with the experimental Hi-C map [6, 1]. The comparisons produced a range of Pearson's correlation coefficients from 0.932 to 0.953. The radial distributions of L-TADs were calculated and the fractions of L-TADs at the NE were compared with the experimental data [15] (selection rule #2 in main text, "Model Development").

To further refine the interaction parameters, in the last stage of model development, we performed simulations of the full ensembles of nuclei topologies (see Fig. 1 in the main text), properly weighted as described in "Methods" (main text), including the use of 3 different ($2.00 \pm 0.06$ $\mu$m) nucleus sizes. As described in "Methods" (main text), the resulting ensemble of simulated nuclei consists of 18 different initial states and the corresponding trajectories; the latter are used to obtain the topology and size averaged chromatin distributions and Hi-C maps for each test combination of the interaction parameters. At this stage, 32 parameter sets have been used to determine the "optimum" WT parameter set and to investigate the effects of parameter deviations onto the chromatin properties. $40 \times 10^6$ time-steps trajectories were generated for each of characterised by a single parameter set. Compared to the previous optimization stage, and based on its results, a more narrow range (3 – 4.5 $kT$) of the LAD-NE interactions parameters was used. Several values of the cross-type $\epsilon_{AB}$ parameter and $\epsilon_A$ parameter (in the range $0.1 - 1.0$ $kT$) were used as well. In addition, 8 sets of the interaction parameters for the Lamin mutant model were considered to investigate its chromatin properties.

After calculating the ensemble averaged Hi-C maps for each parameter set, we selected 9 WT-like sets with the highest (greater than 0.955) Pearson's correlation coefficients with the experimental Hi-C map (selection rule #1 in the main text). Given the range ($1 - 2$ $kT$) used for the interaction parameter $\epsilon_B$, the selection rule #3 is satisfied when the A-B cross-type interaction parameter $\epsilon_{AB}$ does not exceed 0.5 $kT$. This reduces the number of the selected sets from 9 to 5. All these new 5 sets have the same $\epsilon_{AB} = 0.5$ $kT$ and all 3 types of $\epsilon_B$ parameters (for Null, PcG and HP1 beads) equal to 1.5 $kT$, and are characterized by roughly the same Pearson's correlation coefficients, $\sim 0.956$. However, the values of the LAD-NE interaction parameters $\epsilon_L$ are still different (3.0, 3.5 and 4.0 $kT$), allowing us to apply the selection rule #2 for further refinement. Slightly different are the values of the $\epsilon_A$ parameter (0.1 and 0.5 $kT$) and $E_{sp}$ parameter (3.0 and 3.5 $kT$), suggesting that these differences don't have a noticeable effect on the local chromatin structure.

We would like to note here that the Pearson's correlation coefficients for the ensemble averaged model Hi-C maps for these "best" 5 parameter sets are higher than the coefficients for each individual nucleus Hi-C map in the ensemble of nuclei. Most of these single nucleus Hi-C coefficients do not exceed the largest value obtained in the previous parameter optimization stage.

Selecting the A-A interaction parameter $\epsilon_A$ to be 0.1 $kT$, as found in the 4 of 5 sets, and having it lower than $\epsilon_{AB} = 0.5$ $kT$ for the cross-type interactions between different classes of B-type beads (e.g., HP1-PcG interactions), and excluding the only set with $E_{sp} = 3.0$ $kT$ (which has the smallest Pearson's correlation coefficient), we have determined a single set of the parameters (other than $\epsilon_L$) we can use to select the LAD-NE interaction strength $\epsilon_L$ in our WT nucleus model.

To select a single value of the $\epsilon_L$ parameter ($\epsilon = \epsilon_L$ in Eq. 2) that reproduces experimentally observed fraction of LADs near the NE (25% [15], selection rule #2), we analyze the radial distributions of L-TADs in the nuclei models with the selected above set of the interaction parameters. The normalized cumulative distributions shown in Fig. 4 suggest that the optimum value of the L-TAD–NE interaction well depth $\epsilon_L$ is 4.0 $kT$. This value ensures that, on average, 25% of L-TAD bead centers are within 0.09 $\mu$m (average bead radius) from the average position of the NE (2.0 $\mu$m from the nucleus center).

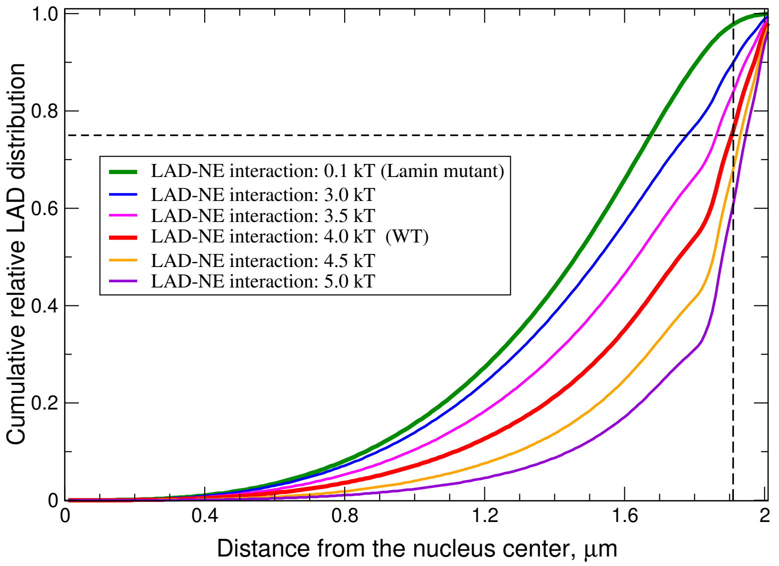


**Figure 4** Cumulative normalized distributions of L-TADs (LAD containing TADs) at different levels of attractive interaction with the nuclear envelope (NE) varying from 0.1 $kT$ to 5 $kT$. The vertical dashed line at R=1.91 $\mu$m marks the center of the bead layer (average bead radius 0.09 $\mu$m) nearest to the NE (with the average internal radius 2 $\mu$m). Experimental data [15] show that about 25% (horizontal dashed line at 0.75 mark) of LADs are on average located at the NE.

The final optimum WT set of the interaction parameters is shown in the second column of the Table 1 in the main text. We use this set for 10x longer simulations reported in the next section. The resulting chromatin density profiles will be compared with the available experimental data [12]. We will also investigate how sensitive is the chromatin structure to the deviations in the parameters.

### Compartmentalization

To see whether our model produces the expected different levels of compartmentalization for A and B types of beads, we used the model derived Hi-C and estimated contact intensities for two types of beads – Active beads (A type) and Null beads (B type). For estimation of the contact intensities we calculated average sums of the contact probabilities for these two types of beads with all other surrounding beads. The resulting values of the sums – 3.86 for Active beads and 5.83 for Null beads – suggest that compartmentalization, characterized by the intensities of contacts, for A type beads is roughly twice less intense than for B type beads.

### Heterogeneity of the biological data used in the model

*Drosophila* nuclei at interphase are modeled using a heterogeneous set of experimental data, which were available to us. The Hi-C data are from late *Drosophila* embryos [6]. The Lamin B mutant data are from larval low-level polytene nuclei that change chromatin distribution similarly to regular diploid nuclei [12, 16]. The LAD data are from *Drosophila* Kc cells of embryonic origin [15, 8]. Since these data are not from highly-specialized tissues, we believe they represent a typical *Drosophila* nucleus. Moreover, LADs appear to be robust in different *D. melanogaster* cells including polytene chromosomes [15].

### Results

#### Temporal evolution of the WT model Hi-C maps

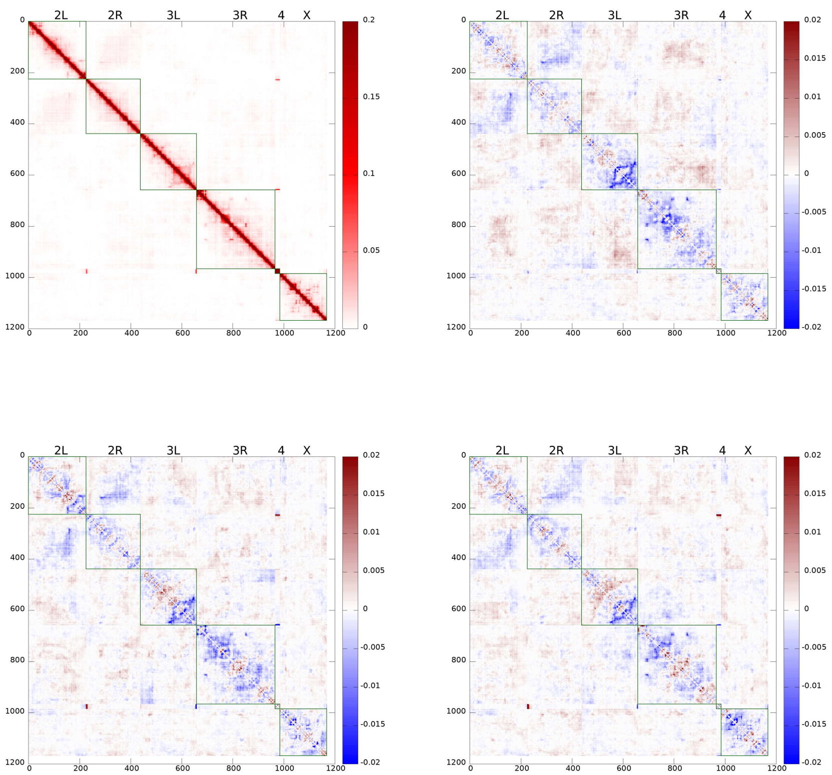


**Figure 5** Details of the temporal evolution of the WT model derived TAD-TAD contact probability (Hi-C) maps. The Hi-C maps are averaged over 5 min ($3 \cdot 10^6$ time steps) time intervals and over the ensemble of 18 system trajectories (see "Methods"). The top left panel is t=0 min Hi-C map, followed (from left to right) by the difference (from t=0 min ) maps at 30 min, 3 hrs and 11 hrs.

#### NE as an "attractive enclosure"

The chromatin distribution in the Lamin mutant nuclei resembles a more condensed, globule-like form compared to WT nuclei. A question arises: does the NE in WT nuclei acts only as an attractive enclosure needed to "keep interphase chromosomes slightly stretched" [16] *on biologically relevant time-scales*? Will the chromatin condense in the complete absence of the NE, not only in the absence of LAD-NE affinity? To answer these questions we have carried out a series of "what if" simulations of the model nuclei with the NE completely removed. A typical result is shown in Fig. 6. The chromosomes partially decondense and extend substantially, and the chromosome chains partially detach from each other resembling an "extended coil" polymer state, on the time scale of a minute. Thus, the NE, even without attractive interactions with LADs, is still necessary to suppress the fluctuations of the chromatin preventing it from unfolding, very quickly.

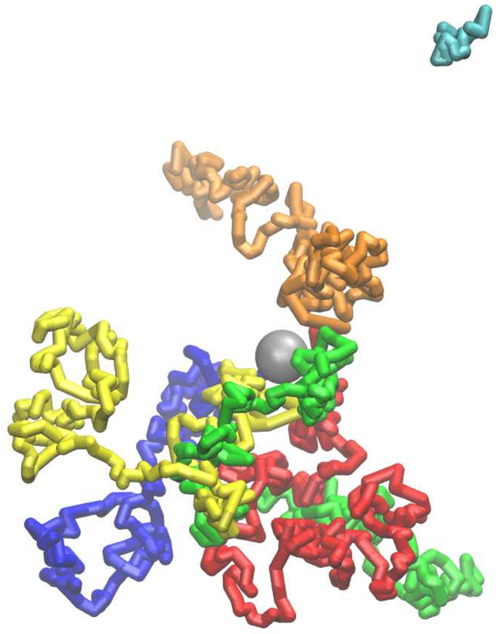

**Figure 6** The effect of complete removal of the NE: chromatin de-compacts. Within 1 minute, chromosome 4 (cyan) dissociates and drifts away from the rest three de-condensed chromosomes.

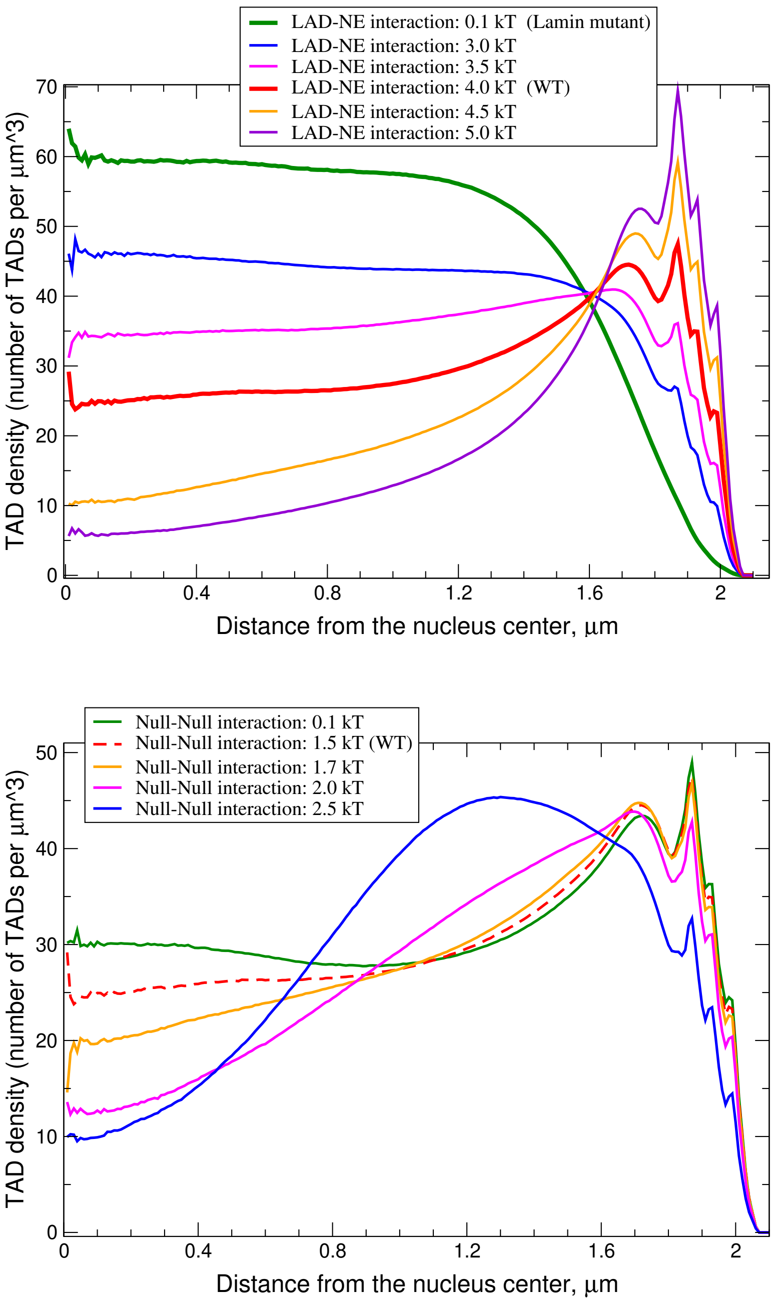


**Figure 7** **Top panel:** Chromatin density distributions in the model nuclei at different levels of LAD-NE attractive interaction (from 0.1 to 5 $kT$). **Bottom panel:** Chromatin density distributions in the model nuclei at different levels of attractive interaction between Null TADs.

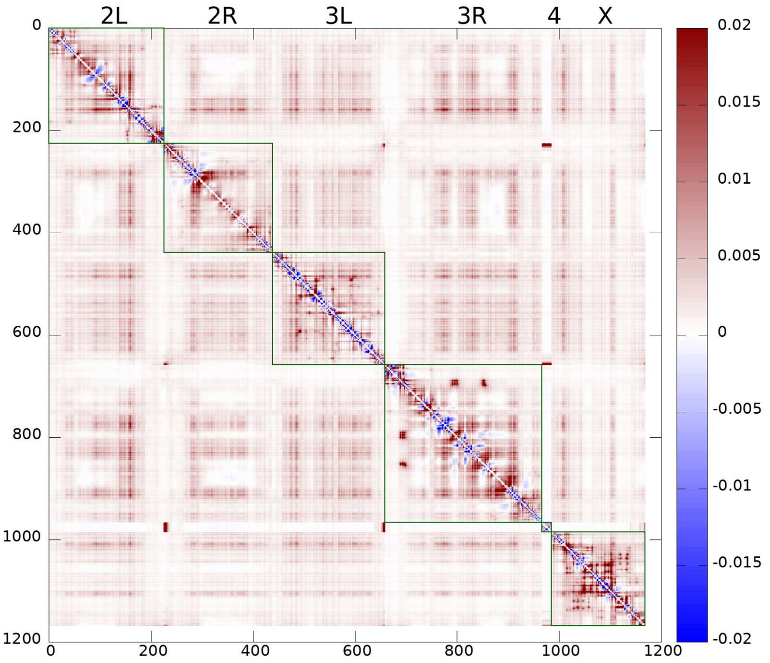


**Figure 8** Difference Hi-C map between Lamin mutant model Hi-C map (Fig. 5, main text) and the corresponding WT model Hi-C map (Fig. 3, main text, bottom panel). Note that the intensity scale is 10 times smaller here than that of the original Hi-C map, pointing to relatively small differences in TAT-TAD contact probabilities. Most TAD-TAD contacts, including inter-chromosome contacts, are slightly enhanced (red areas ) in the Lamin mutant compared to the WT nuclei. Small areas where the contact frequency decreases (blue spots) in the Lamin mutant are limited to close to diagonal intra-arm contacts.

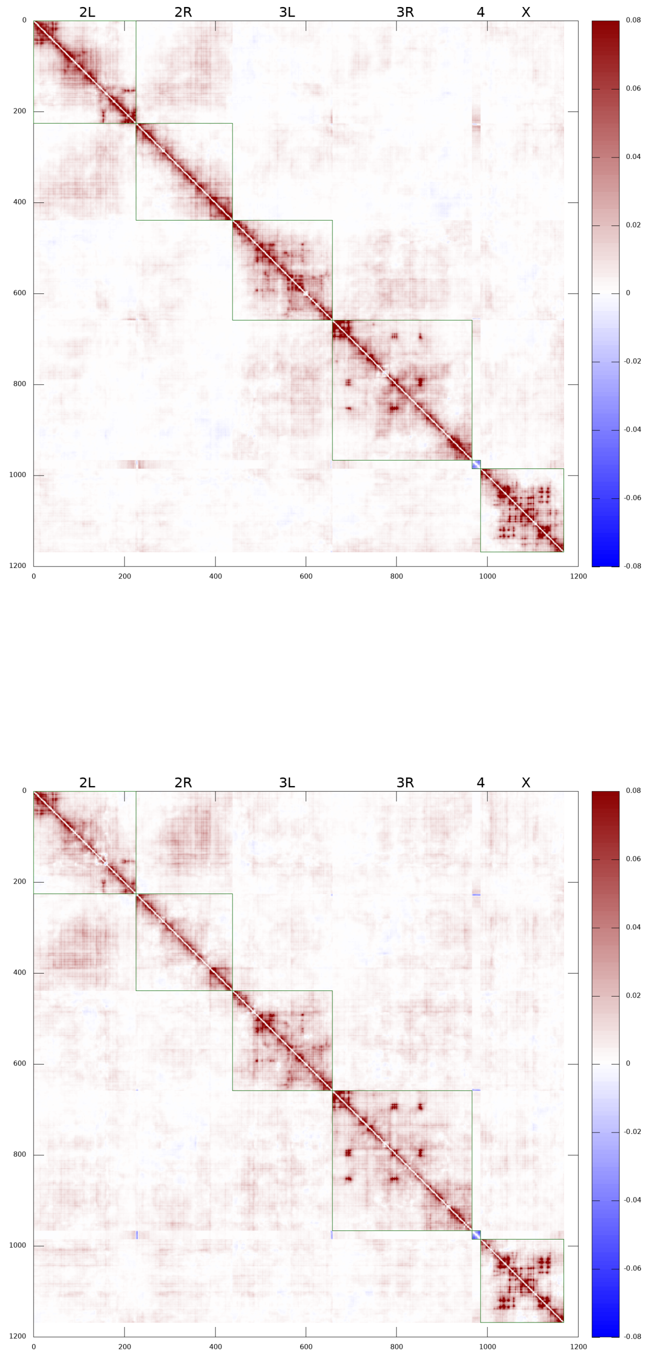


**Figure 9** Increasing the cross-type TAD-TAD interactions (parameter $\epsilon_{AB}$) from 0.5 $kT$ (model selected) to 1.0 $kT$ results in increased TAD-TAD contact probabilities, both in WT and Lamin mutant model nuclei. The Hi-C map differences between modified and selected model nuclei are shown. (**Top panel:**) for the WT model nuclei. (**Bottom panel:**) for the Lamin mutant model nuclei.

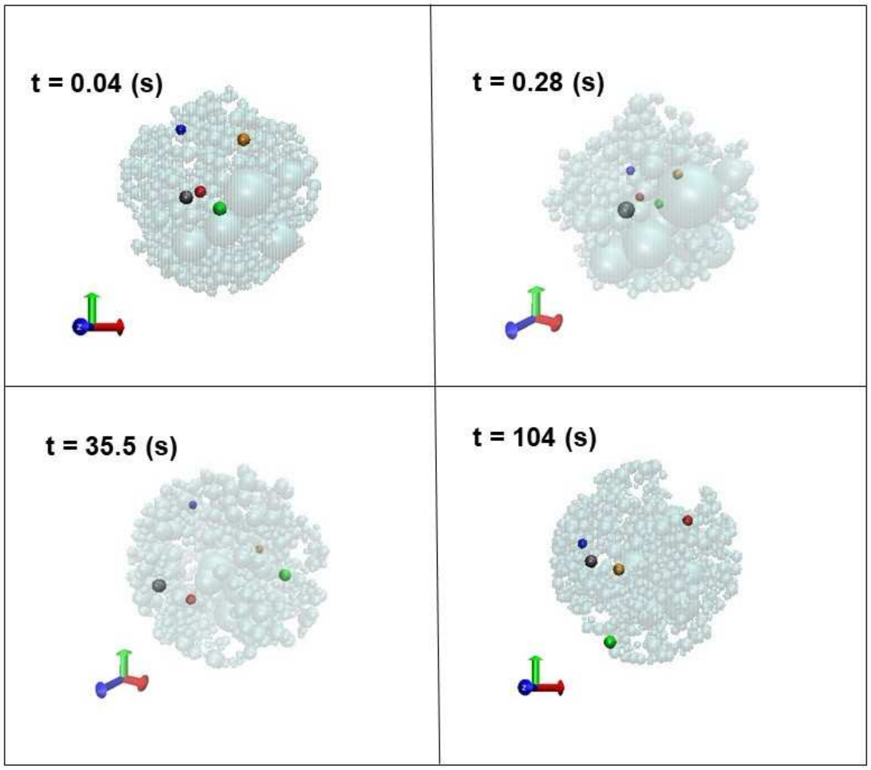


**Figure 10** The mobility of five select L-TADs over about two minutes time interval of the interphase, starting at $t = 0$. Shown are four snapshots from the supplementary movie that represents 20 minutes of time-evolution of model fruit fly nucleus. Each L-TAD is selected from a different chromosome. The L-TADs represented by orange, green, grey, red and blue are from Chr 2L, 2R, 3L, 3R and Chr X, respectively. All other beads, including the large heterochromatic beads, are shown as transparent grey spheres. The upper panel snapshots, from left to right, correspond $t = 0.04$ (s) and $t = 0.28$ (s), respectively. The lower panel snapshots, from left to right, correspond to $t = 35.5$ (s) and $t = 104$ (s), respectively. The coordinate frame vectors are shown in the left corner of each panel. Apparent variation in each L-TAD sphere size conveys the depth perception: spheres closer to the viewer are larger. Image Credit: Samira Mali. Rendering by VMD [17]. Visualization of motion of these L-TADs, from 3 different viewing angles, is available as supplementary movies. (Also at https://www.dropbox.com/sh/11r65jvanyua0re/AAA3QkiS3Ry0BIIa34sCfrtJa?dl=0 ).

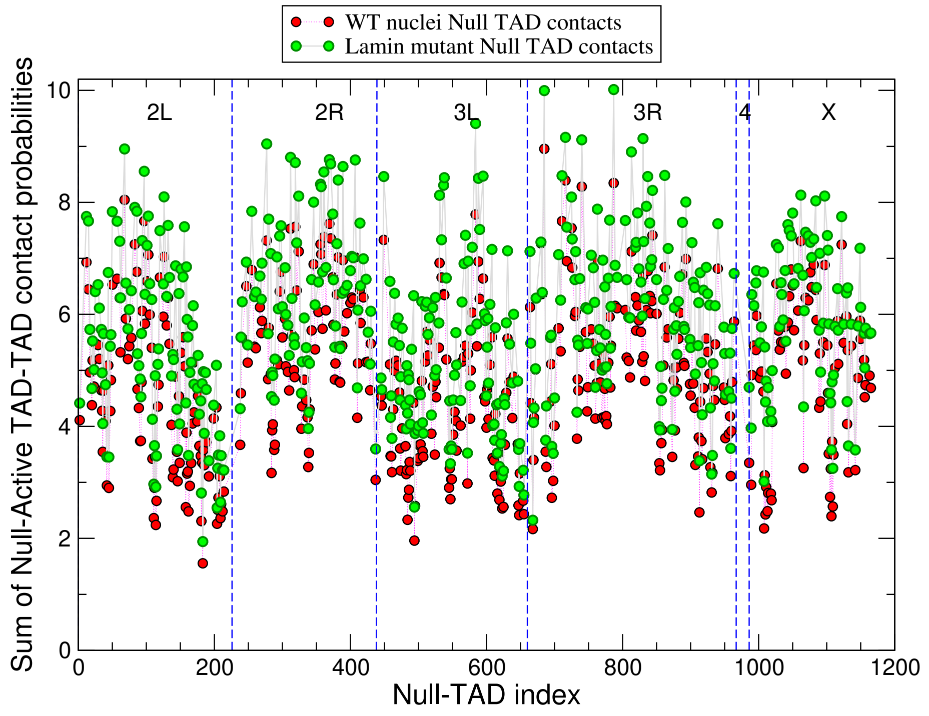


**Figure 11** The sums of contact probabilities of Null TADs (B-type TADs) with Active TADs (A-type TADs) only, calculated from the model Hi-C maps for the WT nuclei (red circles) and Lamin mutant nuclei (green circles), for each Null TAD. The average relative increase of the contact probabilities for Lamin mutant Null TADs is 22%

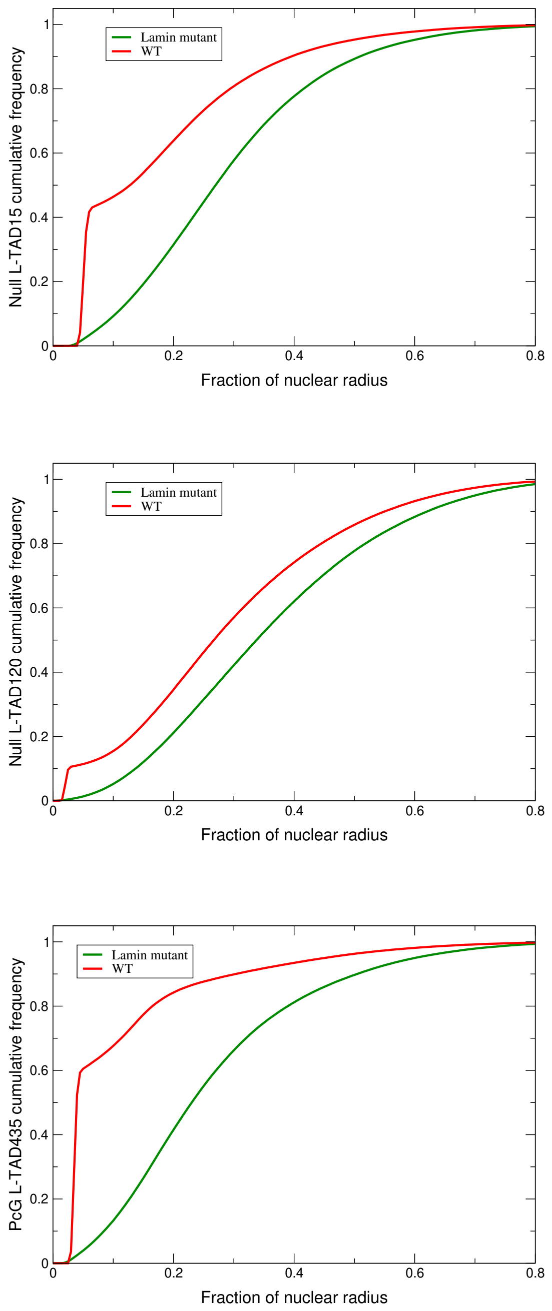


**Figure 12** Predicted cumulative frequencies of radial positions of three cytological regions for which the corresponding experimental data is available. **Top panel:** Cumulative frequencies of radial positions of the Null L-TAD #15 (cytological region $22A$ in Ref. [16]) in the WT and Lamin mutant nucleus models. **Middle panel:** Cumulative frequencies of radial positions of the Null L-TAD #120 (cytological region $36C$ in Ref. [16]) in the WT and Lamin mutant model nuclei. **Bottom panel:** Cumulative frequencies of radial positions of the PcG L-TAD #435 (cytological region $60D$ in Ref. [16]) in the WT and Lamin mutant nucleus models.

**Acknowledgements**
We thank Raju Nadimpalli for help with ESPResSO input scripts, chromatin visualization and Hi-C map processing; Samira Mali for producing the movie clip of the chromosomes dynamics; Yoonjin Kim for technical assistance; Frank Alber for a useful discussion and for providing a comprehensive set of configurational snapshots from Ref. [1]. This study was supported by the National Science Foundation [MCB-1715207].

**Availability of data and material**
The modeling code ESPResSo 3.3.1 [5] used in this research is available at http://espressomd.org/. The software is free, open-source, published under the GNU General Public License (GPL3). The computational protocols for coarse-grained MD simulations and data analysis, as well as data files are available from the authors upon request. Visualization of motion of randomly selected L-TADs in each chromosome on a time-scale of 20 minute is available at https://www.dropbox.com/sh/11r65jvanyua0re/AAA3QkiS3Ry0BIIa34sCfrtJa?dl=0

**Author details**
[1]Department of Computer Science, Virginia Tech, 24061, Blacksburg, VA, USA. [2]Department of Entomology, Virginia Tech, 24061, Blacksburg, VA, USA. [3]Edward Via College of Osteopathic Medicine, 2265 Kraft Drive, 24060, Blacksburg, VA, USA. [4]Department of Physics, Virginia Tech, 24061, Blacksburg, VA, USA. [5]Center for Soft Matter and Biological Physics, Virginia Tech, 24061, Blacksburg, VA, USA.
